# The ciliary kinesin KIF7 controls the development of the cerebral cortex by acting differentially on SHH-signaling in dorsal and ventral forebrain

**DOI:** 10.1101/2024.03.21.586159

**Authors:** María Pedraza, Valentina Grampa, Sophie Scotto-Lomassese, Julien Puech, Aude Muzerelle, Azka Mohammad, Sophie Lebon, Nicolas Renier, Christine Métin, Justine Masson

## Abstract

Mutations of *KIF7*, a key ciliary component of Sonic hedgehog (SHH) pathway, are associated in humans with cerebral cortex malformations and clinical features suggestive of cortical dysfunction. KIF7 regulates the processing of GLI-A and GLI3-R transcription factors in a SHH-dependent manner both in humans and mice. Here, we examine the embryonic cortex development of a mouse model that lacks the expression of KIF7 (*Kif7 -/-)*. The cortex is composed of principal neurons generated locally in the dorsal telencephalon where SHH expression is low and inhibitory interneurons (cIN) generated in the ventral telencephalon where SHH expression is high. We observe a strong impact of *Kif7* deletion on the dorsal cortex development whose abnormalities resemble those of GLI3-R mutants: subplate cells are absent, the intermediate progenitor layer and cortical plate do not segregate properly, and corticofugal axons do not develop timely, leading to a delayed colonization of the telencephalon by thalamo-cortical axons. These structural defects alter the cortical distribution of cIN, which moreover exhibit intrinsic migration defects and cortical trajectories resembling those of cyclopamine-treated cIN. Our results show that *Kif7* deletion impairs the cortex development in multiple ways, exhibiting opposite effects on SHH pathway activity in the developing principal neurons and inhibitory interneurons.

## INTRODUCTION

The primary cilium is a tiny microtubule-based organelle present on the surface of nearly all mammalian cell types including neurons, which functions as a signaling hub and transduces several signaling pathways comprising Sonic Hedghog (SHH), Wnt, Delta/Notch and mTOR pathways ^1^. Primary cilium dysfunction causes pleiotropic diseases named ciliopathies. KIF7 is a ciliary kinesin responsible for the trafficking and the positive and negative regulation of the GLI transcription factors in the primary cilium of mammals ^2^. Loss of function of KIF7 in the mouse has shown that KIF7 regulates SHH signalling by acting downstream of Smoothened (Smo) and upstream of GLI2 and GLI3 ^3–7^. Studies in mice established that KIF7 activity is dependent on the expression level of SHH. In the absence of SHH, KIF7 localizes to the base of the primary cilium ^5^. GLI factors are phosphorylated and addressed to the proteasome at the base of the primary cilium for degradation, leading to the formation of a cleaved and stable transcriptional repressor (GLI3-R). In the presence of SHH, KIF7 accumulates at the distal tip of the primary cilium ^4,5^ and associates with full length GLI2/3 that become transcriptional activators (GLI-A) ^8^.

Since 2011, ten studies have identified patients carrying mutations in the *KIF7* gene responsible of ciliopathies classified according to clinical features as hydrolethalus, acrocallosal, Joubert and Greig cephalopolysyndactyly syndromes ^9–18^. MRI investigations revealed macrocephaly, ventricles enlargement and corpus callosum alterations. They also showed the characteristic hindbrain abnormalities observed in most ciliopathies such as molar tooth sign (MTS) and cerebellar atrophy. Patients also presented with mild but frequent cortical malformations as well as neurodevelopmental delay, intellectual disability and seizures ^11,12,15,19^ which indicate cortical abnormalities.

In physiological conditions, the proper activity of the cortex relies on the excitatory/inhibitory balance i.e. on the ratio, positioning and connectivity of excitatory glutamatergic neurons (principal neurons, PN) and inhibitory GABAergic interneurons (cIN), generated from progenitors in the dorsal and ventral telencephalon, respectively. The cortical neurons of dorsal and ventral origin each migrate towards the developing cortex, the so-called cortical plate (CP), according to a specific timeline allowing the establishment of proper connections in the CP. SHH plays a central role in the forebrain patterning and differentiation. At early embryonic stage, the ventral expression of SHH orchestrates the ventro-dorsal ^20,21^ and medio-lateral ^22^ regionalization of the mouse forebrain and the differentiation of ventral cell types ^20^. *Shh* ablation performed after forebrain patterning alters the specification of distinct subgroups of

GABAergic interneurons ^23–26^. Interestingly, conditional ablation of *Shh* and *Smo* in the embryonic cortex reduces the proliferation of dorsal progenitors ^27^, demonstrating a minimal SHH expression in the dorsal telencephalon, even before birth ^28^. Beside its role on proliferation and specification, SHH also controls the migration of interneurons to the cortical plate ^29–33^. In the embryonic telencephalon, GLI transcription factors mediate SHH signals in complex and specific ways. The three GLI factors identified in mammals are expressed in the mouse forebrain: GLI1 along the source of SHH in the ganglionic sulcus, and GLI2 and GLI3 dorsally to the SHH source ^26,34,35^. GLI1 acts uniquely as a pathway activator whereas GLI2 and GLI3 can be processed in transcriptional activators or inhibitors in the primary cilium. However, both GLI1 and GLI2 function primarily as transcriptional activators in response to SHH activity in the ventral forebrain ^35^. Nevertheless, GLI1 and GLI2 mutants show mild phenotypes ^36,37^ and GLI2 seems required to transduce high level SHH signals in mice ^38^. In contrast, the development of the dorsal cortex, where principal excitatory neurons differentiate, depends mainly on the expression of the GLI3-R repressor, in agreement with the low cortical expression of SHH ^39,40^. Following patterning, Gli3R/A ratio remains critical for specifying the fate of cortical progenitors and regulating cell cycle kinetics ^41–43^.

Previous studies investigated the mechanisms underlying the corpus callosum agenesis in patients with *KIF7* mutation using *Kif7* knock-out (*Kif7* -/-) mice ^44^ and the consequence of KIF7 knock down on cortical neurogenesis in principal cells electroporated with *Kif7* shRNA ^45^. It remains that the influence of developmental abnormalities associated with ciliopathies on the cortical cytoarchitecture is poorly understood. Given the crucial role of KIF7 in regulating the GLI pathways and the important role of Gli3-R and GLI2/3-A in regulating the early developmental stages of the dorsal and ventral telencephalon, respectively, we examined here the cortical development from E12.5, the establishment of long distance projections with the thalamus and the migration of GABAergic interneurons (cIN) in *Kif7* -/- mice. The migratory behavior of *Kif7* -/- cINs was investigated by time-lapse videomicroscopy in co-cultures and in organotypic cortical slices. We also compared the migration of cINs in *Kif7* -/- cortical slices and in control slices treated with pharmacological activator and inhibitor of the SHH pathway. We moreover determined the local distribution of the SHH protein in the embryonic cortex. Our results show developmental defects leading to permanently displaced neurons, abnormal formation of cortical layers and defective cortical circuits that could be responsible for epilepsy and/or intellectual deficiency in patients carrying *KIF7* mutation ^9,11–18,44^.

## RESULTS

### Kif7 knock-out mice as a model to investigate the structural cortical defects that could lead to clinical feature in patients carrying KIF7 mutation

The *KIF7* gene is located on chromosome 15 in human and encodes a 1343 aa protein containing, a kinesin motor domain and a GLI-binding domain in the N-terminal part followed by a Coiled-coil region and a Cargo domain able to bind a diverse set of cargos in the C-terminal part ^46^. Table 1 summarizes the clinical features associated with mutations targeting either the kinesin or GLI binding domain, the Coiled-coil domain, or the N-terminal Cargo domain. Interestingly, various mutations are associated with the same clinical picture, suggesting that *KIF7* mutations, whatever their nature, could lead to protein loss of function, for example by altering the protein structure and solubility as proposed by Klejnot et Kozielski ^47^. All patients carrying mutation in the *KIF7* gene have developmental delay (DD) and intellectual deficit (ID) associated with classical defects of ciliopathies (ventricle enlargement, macrocephaly, corpus callosum agenesis and MTS). Some patients have additional anatomical cerebral cortex defects such as poor frontal development, atrophy, pachygyria and heterotopia ^9,10,13,16^, which could participate not only to DD and ID but moreover to seizure as observed in a quarter of patients. In this context, we used a murine model in which the *Kif7* gene had been deleted (*Kif7* -/-) ^3^ to investigate the consequence of KIF7 loss of function on the cortex development. *Kif7* -/- mice have been previously characterized as dying at birth with severe malformations, skeletal abnormalities (digits and ribs), neural tube patterning defects including exencephaly in a third of the mutants, microphtalmia, lack of olfactory bulbs and CC agenesis ^3,4,44^. In the present study, the KIF7 loss of function was analyzed in a mouse strain with a mixed genetic background in which exencephaly was observed at the same frequency in *Kif7 -/-* and control embryos. As previously reported, *Kif7 -/-* embryos displayed microphtalmia [illustrated at embryonic stage 14.5 (E14.5), Fig. 1A, black arrow] and polydactyly (not illustrated). Interestingly, skin laxity previously reported in ciliopathic patients ^48,49^ was observed in all mutants (Fig. 1A, white arrow). Comparison of brains from *Kif7* -/- embryos and control littermates revealed the absence of olfactory bulbs in *Kif7* -/- brains (illustrated at E14.5, Fig. 1B, white arrows on ventral face view).

**Figure 1.**
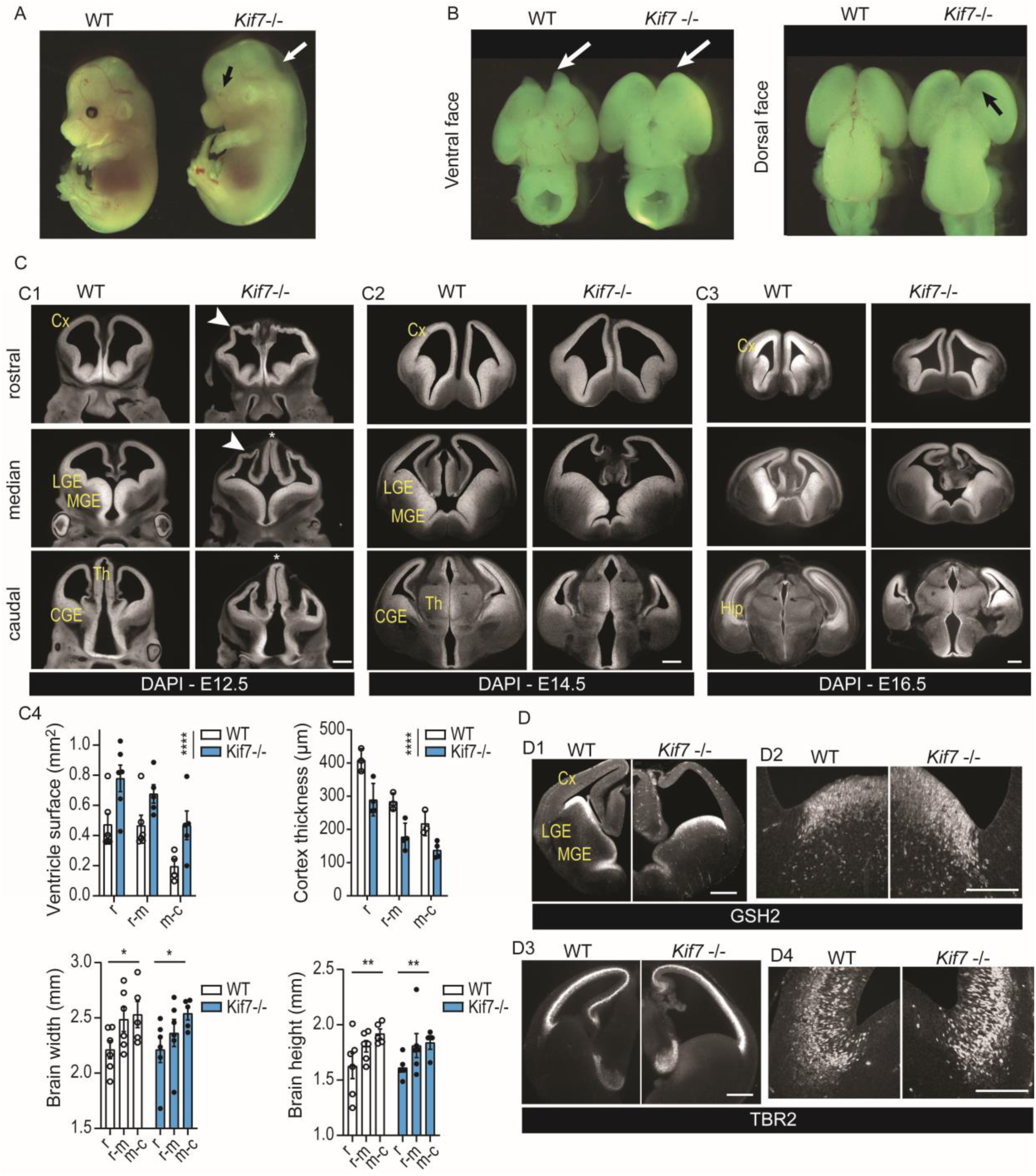
*Kif7* deletion alters cortical development on latero-medial and rostro-caudal axes. **(A)** *Kif7* -/- embryos are microphtalmic (black arrow) and exhibit skin laxity (white arrow). **(B)** External examination of the brain reveals the lack of olfactory bulbs (white arrows, left panel) and the thinning of the dorsal telencephalon (black arrow, right panel). **(C)** DAPI staining of rostro-caudal series of coronal sections at E12.5 (C1), E14.5 (C2) and E16.5 (C3) illustrates the anatomical defects of *Kif7* -/- embryonic brains quantified at E14.5 in C4. The ventricles of *Kif7* -/- embryos are strongly enlarged (upper left graph; WT, n=4-6, *Kif7* -/-, n=5- 6 depending on the rostro-caudal level), their cortical thickness strongly decreased (upper right graph; (WT, n=3; *Kif7* -/-, n= 4), resulting in minimal brain width (lower left graph; WT, n=5- 6; n=5-6 for *Kif7* -/- depending on the rostro-caudal level) and height (lower right graph; WT, n=5-6; *Kif7* -/-, n=4-6 depending on the rostro-caudal level) changes. Statistical significance was tested by Two-way ANOVA or mixed model (GraphPad 8.1.0). For ventricle surface and cortex thickness, mix model reveals a genotype effect (p<0.0001). For brain width and height, no genotype effect was observed, but a significant effect of the rostro-caudal level on brain width (*P*=0.0441) and height (*P*=0.0092). **(D)** The pallium-subpallium boundary identified by the limit of expression of ventral (GSH2, D1, D2) and dorsal (TBR2, D3, D4) telencephalic markers is less precisely defined and slightly shifted to the ventricular angle in E13-E14 *Kif7* - /- embryos compared to wild type embryos (epifluorescent low magnification, D1, D3; confocal high magnification images, D2, D4). Graphs in C4 represent the means and S.E.M. Cx, cortex; Hip, hippocampus; LGE, lateral ganglionic eminence; MGE, median ganglionic eminence; CGE, caudal ganglionic eminence; Th, thalamus.. Scale bars, 500 µm (C,D1,D3), 150 µm (D2,D4).

**Table 1.**
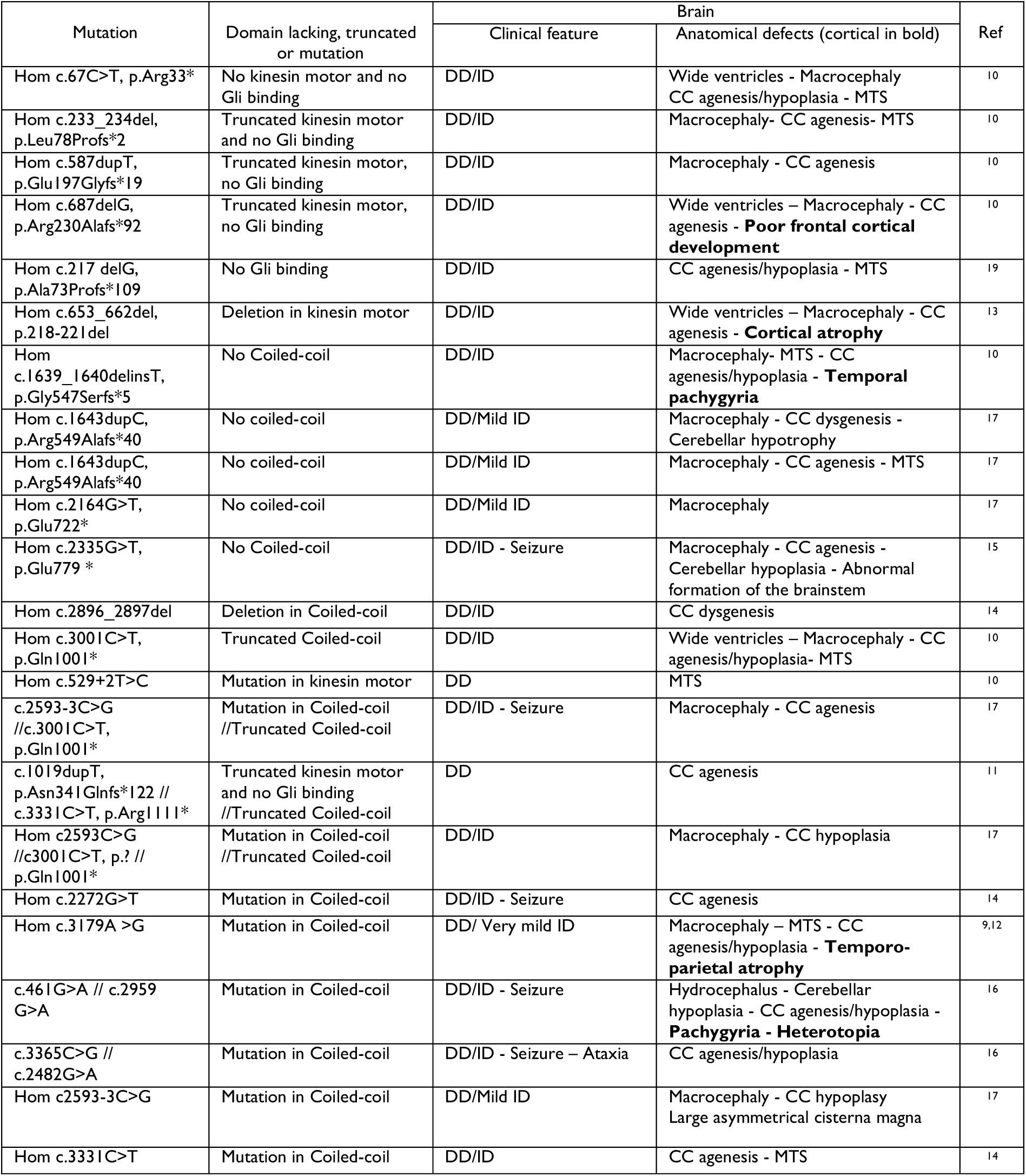
Clinical diagnosis of patients carrying mutation in the *KIF7* gene on both alleles from the literature. The ablated or mutated domains were identified. Clinical features associated with cortical dysfunction are listed. Among cerebral defects, those observed in the cortex are enlighted. DD, developmental delay; ID, intellectual deficit; CC, corpus callosum; MTS, molar tooth sign.

### Kif7 invalidation alters the development of the cortex

The thinning of the dorsal cortex at E14.5 allowing the lateral ventricles to be seen through (Fig. 1B, black arrow on dorsal face view) proned us to investigate the structural organization of the forebrain over time (Fig. 1C1-3, E12.5-E16.5). Analyses performed on rostro-caudal series of frontal sections identified strong abnormalities in the cortex development. E12.5 *Kif7* -/- embryos exhibited cortical wall folding and thinning in rostro-median sections (Fig. 1C1, arrowheads) and prominent diencephalon (Fig. 1C1, stars). At E14.5, cortex folding was not longer observed in *Kif7 -/-* embryos which exhibited enlarged lateral ventricles (compare left and right panels in Fig.1C2). The cortex remained thinned in the dorsal region and the latero- dorsal extension of the telencephalic vesicle was strongly reduced caudally. These defects were maintained in E16.5 embryos (Fig. 1C3) and the hippocampus did not develop properly (Fig. 1C3, caudal section). A quantitative analysis performed in E14.5 brains (Fig.1C4) confirmed the ventriculomegaly (upper left graph) and the cortex thinning (upper right graph) in the rostral and median sections despite unchanged total width and height of the telencephalon (Fig. 1C4, lowers graphs). KIF7 depletion in E10.5-E11.5 mouse embryos was reported to alter the dorso-ventral patterning of the medial forebrain ^44^. The frontier of expression of GSH2 (Fig. 1D1,D2) and TBR2 (Fig. 1D3,D4), two transcription factors expressed in the ventral and dorsal telencephalon respectively, was slightly abnormal at E13-E14 in *Kif7* -/- embryos with a small shift of GSH2(+) cells toward the ventricular angle (Fig. 1D2) and a dispersion of some TBR2(+) cells at the pallium-subpallium boundary (PSB) (Fig. 1D4). This suggests a near-normal dorso-ventral patterning of the telencephalic vesicles, however with a discrete blurring of the dorsal and ventral markers at the PSB.

In *Kif7* -/- embryos, the cleavage of full length GLI3 (GLI3-FL) into the transcriptional repressor GLI3 (GLI3-R) is decreased ^3,4^. Here, we analyzed the expression of GLI3-FL and GLI3-R in the cortex and in the medial ganglionic eminence (MGE) where cortical interneurons (cIN) are born. We showed minimal expression of GLI3-R and GLI3-FL in the MGE compared to the cortex (Fig. 2A) and a strong decrease in the cleavage of GLI3-FL in GLI3-R only in the cortex of *Kif7* -/- embryos (Fig. 2A,B).

**Figure 2.**
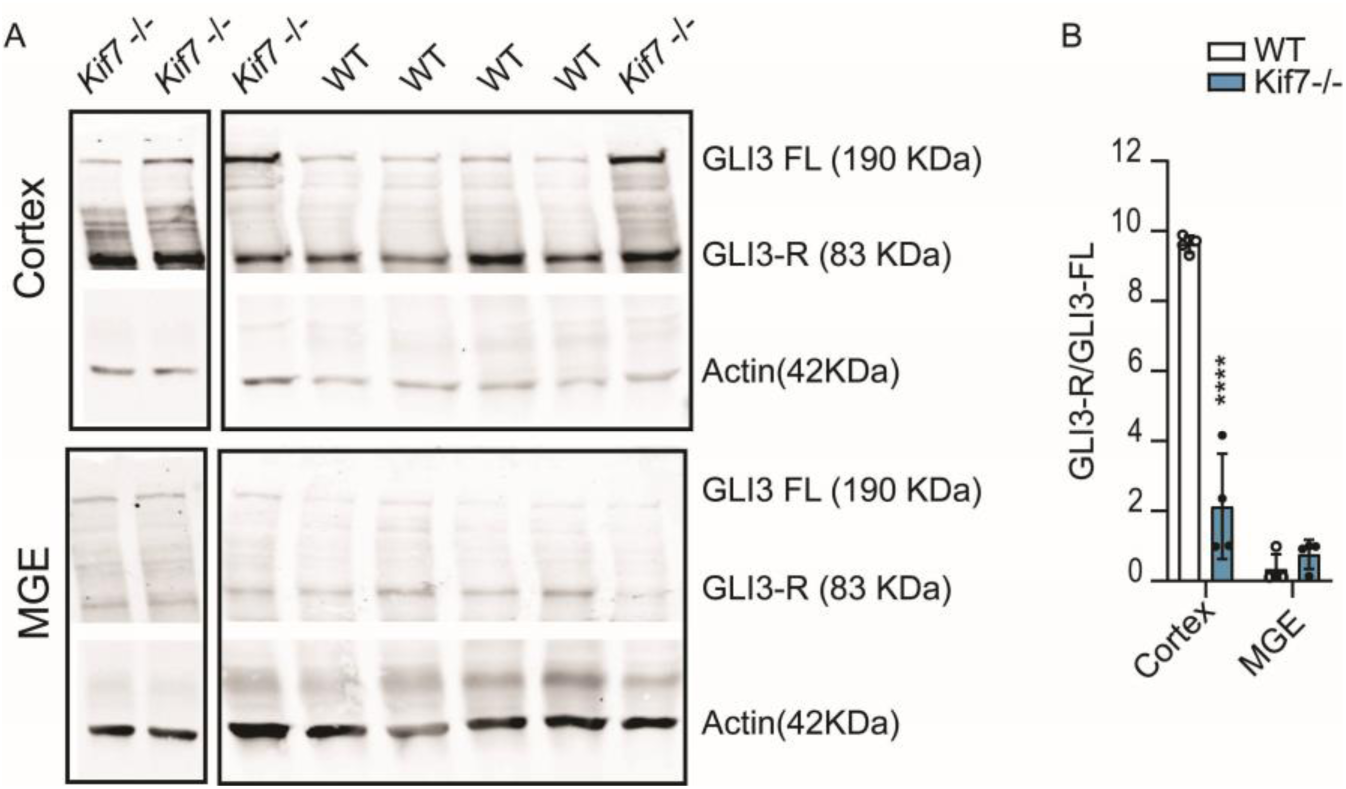
Western blot analysis on the cortex and MGE of wild type (WT) and *Kif7* -/- embryos at E14.5. **(A)** The cleaved form of GLI3 (GLI3-R at 83 KDa) and the full length GLI3 (GLI3-FL at 190 KDa) are more abundant in the cortex compared to the MGE (see actin band intensity for protein loading). **(B)** The ratio Gli3-R/Gli3-FL is lower in the MGE than in the cortex of WT animals and is significantly decreased only in the cortex of *Kif7* -/- brains compared to control. Two way ANOVA reveals significant interaction between brain structure and genotype (WT, n=4; *Kif7 -/-*, n=4; *P*=0.002) and multiple comparisons show a statistical difference between genotype only in the cortex (****, *P*<0.0001)]. Graph represents the means and S.E.M.

The structural defects described above convinced us to examine the cellular organization of the cortex of *Kif7* -/- embryos. At E12.5, the whole cortex is proliferative. From stage E14.5, the mouse cortex can be described as a stack of three specialized domains: i) the upper/superficial cortical layers containing post-mitotic cells generated in the proliferative zones of the cortex, with TBR1(+) post-mitotic neurons located in the cortical plate (CP) in-between the subplate (SP) and marginal zone (MZ) MAP2-expressing cells; ii) the deep proliferative layers, ventricular and subventricular zones (VZ and SVZ, respectively), located along the lateral ventricle; and iii) the intermediate zone (IZ) located between the CP and the proliferative layers, which hosts the radially and tangentially migrating neurons and growing axons. At the end of the embryonic development, the proliferative layers are strongly reduced due to neurogenesis decrease, whereas the cortical thickness had increased due to CP delamination. At E14.5, the TBR1(+) CP cells appeared more clustered in *Kif7 -/-* embryos than in wild type (WT) embryos where they formed a rather regular, radially organized CP (compare left and right panels in Fig. 3A1,3A2). The thickness of the upper MAP2(+) MZ layer was extremly irregular in *Kif7 -/-* embryos and MAP2(+) SP cells were missing in the dorsal cortex (Fig. 3A1,A2, white arrows indicate the limit of distribution of MAP2(+) SP cells). In E16.5 and E18.5 embryos, the CP was obviously thinner in the mutant as compared to the WT (Fig. 3B,C), even though CTIP(+) cells differentiated normally in V-VI layers (Fig. 3C). The CP thinning appeared even more pronounced in the caudal than in the median and rostral cortex. We then analyzed the proliferative layers focusing on TBR2(+) intermediate progenitors (IP) which normally form a dense layer in the cortical SVZ (Fig. 3D1, left). The TBR2(+) layer constantly showed an abnormal positioning in the dorsal cortex of *Kif7* -/- embryos. TBR2(+) cells indeed reached the brain surface (Fig. 3D1, arrow) where they mixed up with post-mitotic TBR1(+) cells instead of keeping a minimal distance with TBR1 cells as observed in control brains (Fig. 3D2, arrow). As a consequence intermediate progenitors and post-mitotic neurons no longer segregated in the dorsal cortex of *Kif7* -/- embryos where no IZ was observed (Fig. 3D2, arrow). Accordingly to the thinning of the cortex starting at E12.5, the TBR2(+) layer was thinner at all development stages analyzed (Fig. 3E,F). In contrast, the abnormal positioning of TBR2(+) cells at the brain surface in the dorsal cortex of E14.5 embryos was no longer observed at E16.5 (Fig. 3F) suggesting either a transient structural disorganization and/or developmental delay of the dorsal cortex in Kif7-/- embryos at E14.5. The TBR2 staining moreover revealed focal heterotopia in the dorsal or lateral cortex of E14.5 *Kif7* -/- embryos, either at the ventricular or pial surface (Fig. S1).

**Figure 3.**
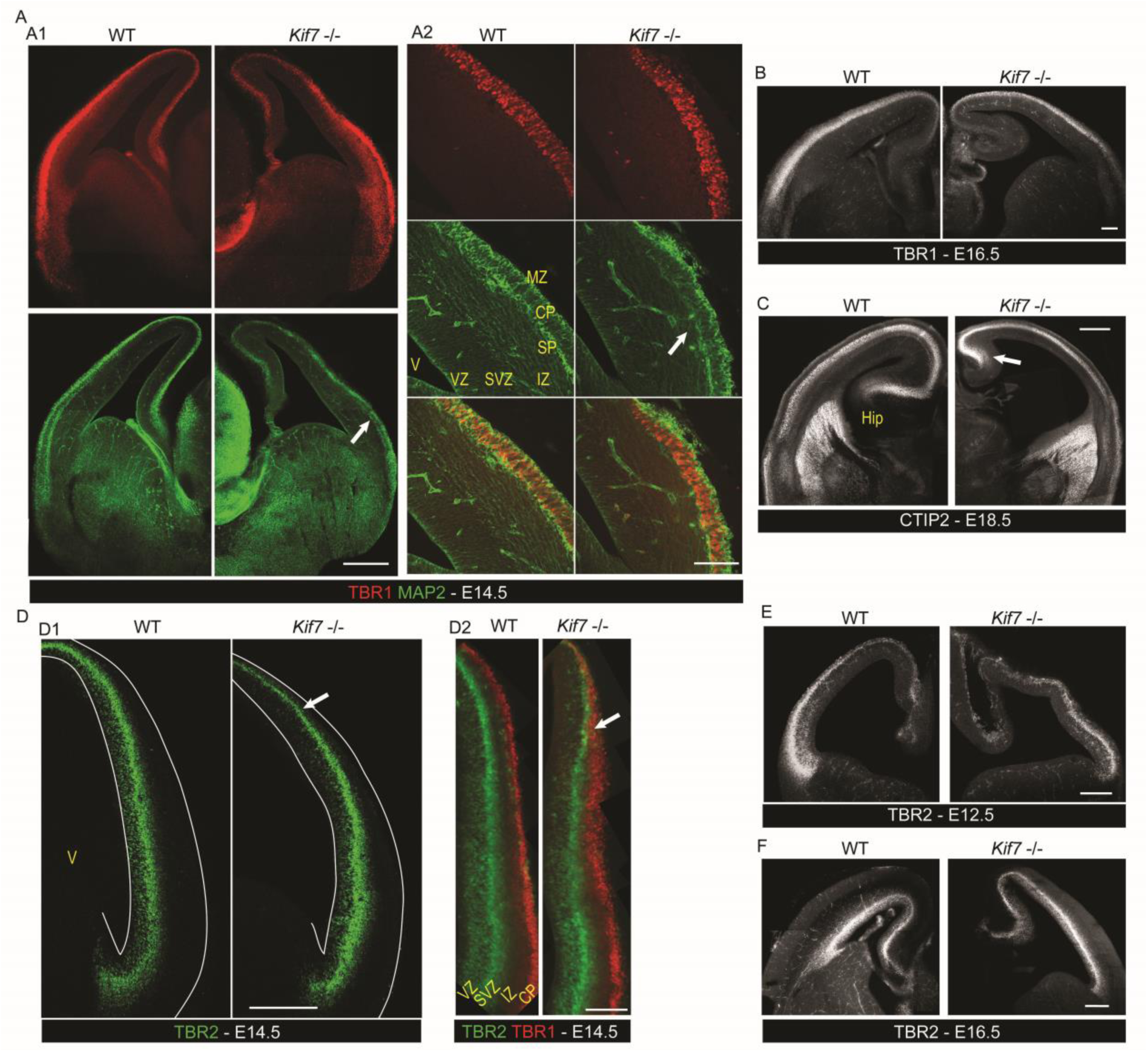
Histological alterations in the developing cortex of *Kif7* -/- embryos. **(A-C)** Immunostaining of cortical post-mitotic layers in E14.5 (A), E16.5 (B) and E18.5 (C) in median coronal sections representative of 5 WT and *Kif7* -/- embryos imaged on epifluorescence (A1,B,C) or confocal (A2) microscopes. At E14.5 (A1,A2), the TBR1(+) staining (red) of the cortical plate is more clustered in *Kif7* -/- than in WT embryos, and the MAP2(+) staining (green) of the subplate is absent in the dorsal cortex of E14.5 *Kif7* -/- embryos (white arrow, right column). The post-mitotic layers remain thinner in *Kif7* -/- embryos at later embryonic stages as illustrated by TBR1 staining at E16.5 (B) and CTIP1 staining at E18.5 (C) that specifically labels the deeper cortical layers (V-VI). Moreover, the hippocampus is underdeveloped in the mutant (white arrow) (C). **(D-F)** Immunostaining of TBR2(+) proliferative layer. At E14.5 (D1,D2), the TBR2(+) layer (green) of secondary progenitors appears disorganized in the lateral cortex of the *Kif7* -/- embryos (white arrowhead in D1) and reaches the brain surface in the dorsal cortex of *Kif7* -/- embryo where it intermingles with post-mitotic TBR1(+) cells (D2, red) (white arrows). In E12.5 *Kif7* -/- embryo (E), the TBR2(+) layer is thin and clustered however its localization in the thickness of the cortex is normal. At E16.5 stage (F), the TBR2(+) layer is normally positionned in the deeper layer of the cortex in both WT and *Kif7* -/- embryos. Hip, hippocampus ; V, ventricle; VZ, ventricular zone; SVZ, subventricular zone; IZ, intermediate zone; CP, cortical plate ; MZ, marginal zone. Scale bars: 250 µm (A1,B,F), 100 µm (A2), 200 µm (D,E); 500 µm (C).

### The loss of Kif7 alters the connectivity between the cortex and the thalamus

The intermediate zone (IZ) hosts migrating neurons and the growing cortical and thalamic projections ^50^. After reaching the CP by radial migration, post-mitotic neurons extended pioneer axons oriented tangentially in the IZ and directed to the PSB ^51,52^. We thus examined whether the structural defects of the CP and SP and the lack of IZ in the dorsal cortex of *Kif7* -/- embryos did associate with developmental abnormalities of corticofugal projections. We labeled corticofugal axons by inserting small crystals of DiI in the CP of E14.5 paraformaldehyde fixed brains. After DiI had diffused along corticofugal axons, we analyzed labeled axons on coronal sections. In both control and *Kif7* -/- brains, corticofugal axons extended below the CP toward the PSB (Fig. 4A). According to the latero-medial gradient of cortical development, DiI injections in the dorsal cortex of control and *Kif7* -/- brains (Fig. 4A1) labeled much less axons than injections in the lateral cortex (Fig. 4A2,A3). Remarkably, the cortical bundles labeled from similar cortical regions were always much smaller and shorter in *Kif7* -/- than in WT brains (compare right and left columns in Fig. 4A). Dorsal injections labeled large bundles crossing the PSB in WT brains, but only a few axons reached the PSB in *Kif7* -/- brains (Fig. 4A1). The growth of corticofugal axons appeared severely hampered in the dorsal cortex of *Kif7* -/- embryos. Lateral injections in WT brains labeled anterogradely corticofugal axons that reached the embryonic striatum and continued medially in the internal capsule (IC), and labeled retrogradely thalamic projections (Fig. 4A3, left panel). Lateral injections in *Kif7* -/- embryos labeled anterogradely cortical axons that spread in the striatum (Fig. 4A2, right panel) and labeled a large bundle directed to the ventral pre-optic area (Fig. 4A3, right panel, white arrow), which was never observed in WT brains. Moreover, no thalamic axons were retrogradely labeled from the lateral cortex in *Kif7* -/- embryos.

**Figure 4.**
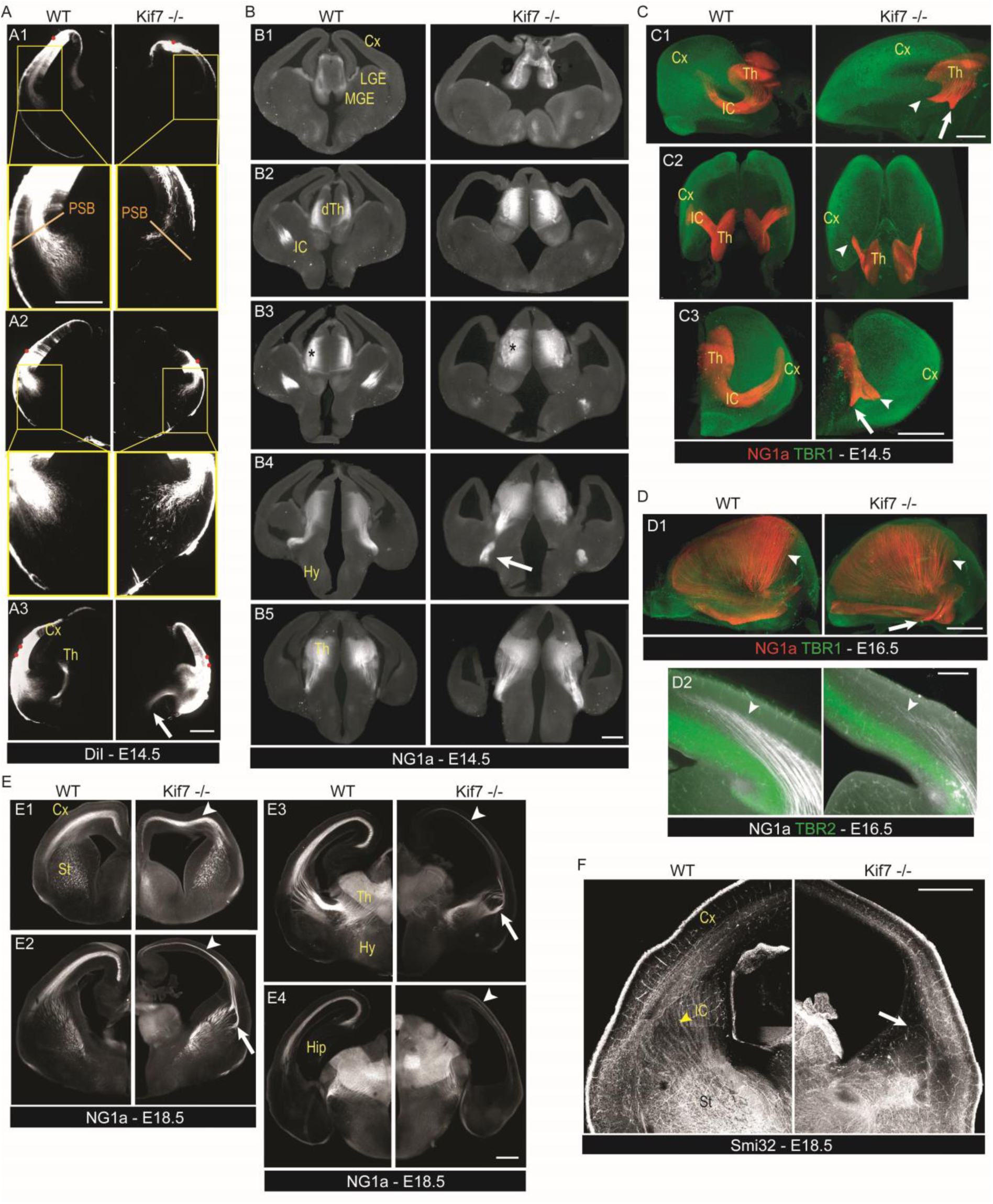
*Kif7* deletion disrupts of the connectivity between the cortex and the thalamus. **(A)** Panels illustrate the corticofugal projections labeled by DiI crystal (red dots) positioned in the dorsal (A1) and lateral (A2,A3) cortex of E14.5 WT (left) and *Kif7* -/- (right) embryos on vibratome sections performed 30 days after DiI placement and imaged on a macroscope. In *Kif7 -/-* embryos, less axons project from the dorsal (A1) and lateral (A2) cortex to the subpallium than in WT (compare enlarged views of the projections below A1 and A2). Cortical injections in *Kif7 -/-* embryos do not label thalamic axons (compare left and right panels in A3) but label a ventral projection (A3, white arrow). **(B)** Rostro-caudal series of coronal sections (B1-B5) immunostained with anti-Netrin G1a (NG1a) antibodies compare the trajectory of thalamo-cortical axons (TCA) in a E14.5 WT (left) and in a *Kif7 -/-* (right) embryo imaged on a macroscope. TCA reach the pallium-subpallium boundary of the WT embryo whereas they are lost in the ventral forebrain of the mutant (B4, white arrow). **(C)** Representative three-dimensional reconstructions (C1, lateral; C2, horizontal; C3, coronal views) of WT and *Kif7 -/-* E14.5 brains immunostained as a whole with NG1a (red) and TBR1 (green) antibodies that label respectively the TCA and cortical plate cells before transparization and imaging with a light sheet microscope (see Supplementary movies S1 and S2). While all labeled TCA extend in the internal capsule (IC) and a significant proportion of them enter the cerebral cortex in the WT brain, TCA in the *Kif7* -/- brain split in two bundles in the basal telencephalon. A bundle stops shortly after entering the IC (white arrowheads) whereas the second bundle extends ventrally (white arrows). **(D)** At E16.5, TCA are immunostained with NG1a in WT (left) and *Kif7 -/-* (right) embryos in whole brain with TBR1 antibodies before transparization and imaging (D1) and on coronal section with TBR2 (D2). The TCA extend in the cortex in *Kif7* -/- brain, however, fiber density is reduced in the median and caudal brain compared to WT (D1, white arrowhead). TCA invade the post mitotic layers in the cortex (above TBR2 layer) to a lower extend in *Kif7* -/- brain (D2). In the *Kif7* -/- brain, thick bundles of NG1a(+) fibers project ventrally from the caudal telencephalon, a projection never observed in control brains (D1, white arrow). **(E)** At E18.5, rostro-caudal series of coronal sections (E1-E4) immunostained with NG1a antibodies compare the trajectory of TCA in WT (left) and *Kif7 -/-* (right) embryos imaged on a macroscope. TCA reach the dorsal cortex in both WT and *Kif7 -/-* embryos. However, the TCA projection in *Kif7* -/- (arrowheads) become thinner in median sections (E2) and almost disappers in caudal sections (E3,E4). Fiber trajectories in the lateral striatum are abnormal (E2,E3, arrows). **(F)** Panels illustrate the cortical projections immunolabeled by Smi32 antibodies in E18.5 WT and *Kif7* -/- brain. In *Kif7* -/- brain, the number of fibers connecting the cortex with other brain structures is strongly reduced (arrow). Cx, cortex; dTh, dorsal thalamus; Hip, hippocampus; Hy, hypothalamus; St, striatum; IC, internal capsule; LGE, lateral ganglionic eminence; MGE, median ganglionic eminence; PSB, pallium-subpallium boundary; .. Scale bars: 250 µm (A,B,C,D), 500 µm (E,F).

To identify the trajectories of thalamic axons in *Kif7* -/- embryos, we immunostained coronal sections of E14.5 brains with antibodies against the Netrin G1a (NG1a), an early marker of thalamocortical axons (TCA, Fig. 4B) ^53^. At E14.5, NG1a antibodies labeled a larger population of neurons in the dorsal thalamus of *Kif7* -/- than in WT brains (Fig. 4B, stars). In control brains, TCA made a right angle turn to join the IC in the striatum. Then they extended to the PSB and some axons entered the lateral cortex (Fig. 4B2,B3, left panel). In *Kif7* -/- embryos, most thalamic axons formed a thick bundle oriented ventrally in the basal forebrain (Fig. 4B4, white arrow in right panel). A minor structural defect was evidenced at the telo-diencephalic junction by Pax6 staining in *Kif7* -/- brains (Fig. S2), recalling the prethalamus defect described in the ciliary Rfx3 mutant ^54^. Given the complex trajectory of NG1a thalamic axons, we immunostained thalamic axons in whole brains and imaged them after transparisation using a light-sheet microscope. The three-dimensional reconstruction of labeled projections confirmed that thalamic axons made a sharp turn to reach the IC and then navigated straight to the PBS in WT brains (Fig. 4C, left panels and Movie S1). In *Kif7* -/- embryos, most thalamic axons stopped their course after leaving the diencephalon, and formed two short bundles: a large one oriented to the IC (Fig. 4C, right panels, arrowheads and Movie S2) and another one oriented to the amygdala, more caudally and ventrally (Fig.4C, white arrows). This last projection was never observed in WT brains. Similar analyses performed at E16.5 showed that two days later, most TCA had reached the cortex in both WT and *Kif7* -/- embryos (Fig. 4D, arrowheads) despite a small delay in *Kif7* -/- embryos and abnormal trajectories in the PSB region (Fig. 4D1,D2). The ectopic ventral caudal projection to the amygdala was still present at E16.5 in *Kif7* -/- embryos (Fig.4D1, arrow). At E18.5, the TCA projections in the *Kif7* -/- embryos appeared almost normal in the rostral cortex (Fig. 4E1) but drastically reduced in the median and caudal cortex (Fig. 4E2, E3, E4, arrowheads). In addition, a thick bundle of thalamic axons followed an abnormal trajectory at the PSB (Fig. 4E2,E3, arrow). Neurofilament staining with Smi32 antibodies confirmed the strong reduction of projections in the cortex and IC of *Kif7 -/-* embryos (Fig. 4F).

### Kif7 invalidation alters the cortical distribution of cIN at E14.5

Because cIN are born in the basal forebrain and likely depend on contact/functional interactions with pioneer corticofugal projections for the first stages of their migration ^55,56^, we examined their distribution in the developing cortex of *Kif7* -/- animals. cIN are generated outside of the cortex, in the medial and caudal ganglonic eminences (MGE, CGE) and preoptic area (POA) of the ventral forebrain. They enter the lateral cortex at E12.5 - E13.5, depending on mouse strains, and colonize the whole cortex by organizing two main tangential migratory streams, a superficial one in the MZ, and a deep and large one in the lower TBR2(+) IZ/SVZ ^57,58^. We thus analyzed the cortical distribution of tdTomato(+) MGE-derived cIN in WT and *Kif7* -/- Nkx2.1- Cre;Rosa26-tdTomato transgenic embryos. An abnormal MGE cell distribution was observed in the developing cortex of *Kif7* -/- embryos from E14.5 (Fig. 5 and Fig.S3). In mutant brains, both the latero-medial extent of the superficial and deep tangential migratory streams and the space separating them were significantly shortened compared to WT brains (Fig. 5A2,B2, B3). The closeness of the two streams in mutants was consistent with the reduced thickness of the corticofugal and TCA projections that navigate tangentially in the IZ (illustrated in Fig. 4). The superficial migratory stream was thinner and denser in *Kif7 -/-* embryos compared to WT (Fig. 5A1,5B1, right panels). Remarkably, cIN no longer colonized the dorsal cortex of *Kif7* -/- embryos and the deep tangential stream stopped in the region where the layer of TBR2(+) SVZ cells switched to the cortical surface (Fig. 5A1, white arrow). The tangential progression of cIN in the developing cortex is controlled by CXCL12, a chemokine transiently expressed in TBR2(+) cells and in meninges ^60,61^. CXCL12 prevents the premature occurrence of the tangential to radial migration switch required for cIN to colonize the CP ^59–62^. *In situ* hybridization experiments revealed that *CxCl12* mRNA was not expressed in the dorsal cortex of E14.5 *Kif7* -/- embryos (Fig. 5C, black arrow). To characterize the consequences of the abnormal expression pattern of CXCL12 at E14.5 on the migratory behavior of cIN, we sliced coronally E14.5 *Kif7 -/-* and WT forebrains and imaged by time-lapse video-microscopy the trajectories of MGE-derived tdTomato-expressing cIN in the dorsal cortex of organotypic slices (Fig.5D). While most cIN presented tangential and oblique trajectories in WT slices, a majority of cIN exhibited radially oriented trajectories in *Kif7* -/- slices (Fig.5D). CXCL12 expression in SVZ strongly decreases at E16.5 ^59,63^ and analyses performed at E16.5 in fixed *Kif 7* -/- and WT embryos showed that cIN had pursued their migration and reached the dorsal cortex (Fig.S3A) even though the tangential migration of cIN in *Kif7* -/- embryos remained delayed as compared to WT embryos. Analyses performed in a surviving P0 animal (Fig. S3B,C) revealed an abnormal distribution of cIN in the CP characterized by a strong decrease of cIN density in the infragranular layers that were moreover much thinner than in control brains (see CTIP2 staining in Fig. S3C).

**Figure 5.**
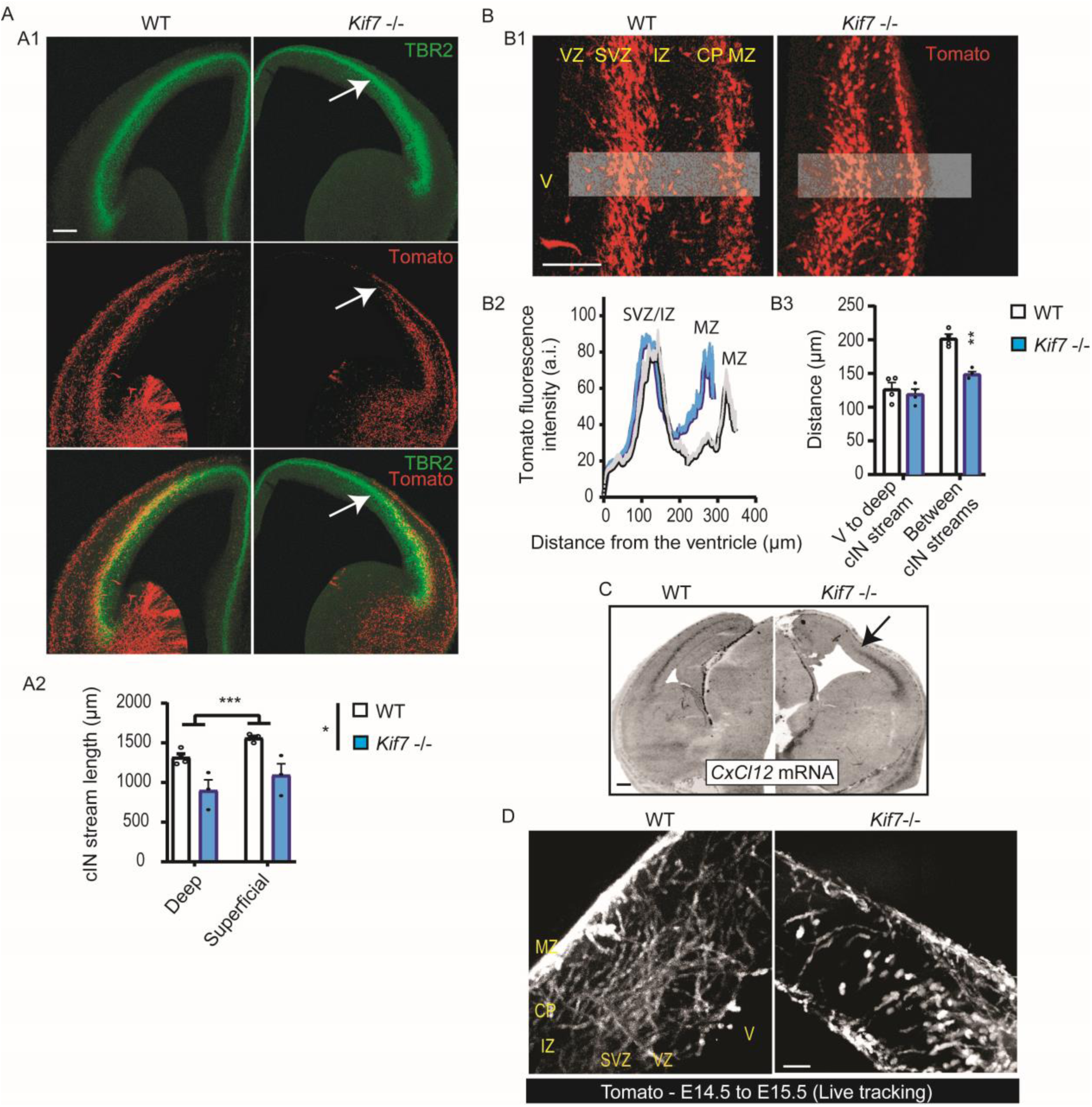
Abnormal cortical distribution of cIN and *Cxcl12* transcript expression in E14.5 in *Kif7 -/-* brains. **(A)** The cortical distribution of cIN is visualized in WT and *Kif7* -/- mouse embryos crossed with the Nkx2.1-Cre/R26R-tdTomato strain in which MGE-derived cIN express the fluorescent marker tdTomato. Panels in A1 compare the distribution of tdTomato (+) cIN in WT and *Kif7* -/- cortical sections prepared at the same rostro-caudal level and in which SVZ is immunostained with TBR2 antibodies (green). Pictures show that the deep migratory stream of cIN terminates in *Kif7* -/- brains in the cortical region where the TBR2(+) layer reaches the cortical surface. Quantitative analysis of the mean length of the deep and superficial migratory streams measured from the entry in the pallium to the last detected cIN in the cortex is illustrated in graph A2. Statistical significance is assessed using Two way ANOVA [layers (WT, n=4; *Kif7 -/-*, n=3; ***, *P*=0.001) and genotype (*, *P*=0.0157)]. **(B)** Representative pictures (B1) of the deep and superficial tangential migratory streams of cIN in the lateral cortex of WT and *Kif7* -/- embryos. Pictures illustrate the decreased thickness of the superficial stream, and the reduced distance between the deep-superficial streams in *Kif7* -/- embryos. Graph in B2 (WT, n=4; *Kif7 -/-*, n=4) compares the distribution of the fluorescence intensity along a ventricle/MZ axis (see grey rectangles in B1) using the plot profile function of FIJI. Curves show no change in the distance between the ventricular wall and the deep cIN, but a significant reduction of the distance between the two streams in the *Kif7* -/- cortical sections as quantified on the graph B3 [WT, n=4; *Kif7 -/-*, n=4; Two way ANOVA reveals a significant interaction between genotype and layer (*P*=0.0233) and multiple comparisons, a statistical difference between genotype only for the distance between de cIN streams (**, *P*=0.0051)]. **(C)** Panels compare the distribution of *Cxcl12* mRNA in WT (left panel) and *Kif7* -/- (right panel) forebrain coronal sections at E14.5. The WT section shows *Cxcl12* transcript enrichment in a deep cortical layer already identified as the SVZ. In the *Kif7* -/- cortical section, the expression of *Cxcl12* transcripts is reduced to the lateral part of the SVZ. (D) Z-projections of 30 frames acquired during 12h in the dorsal cortex of living organoptypic forebrain slices representative of E14.5 WT and *Kif7* -/- embryos with tdTomato expressing cIN. *Kif7* -/- cIN were able to migrate dorsally but followed preferentially radially oriented trajectories Scale bars: 200 µm.

The abnormal distribution of cIN in the cortex of *Kif7 -/-* embryos reflected the structural and molecular abnormalities that we had identified in their developing cortex. We next examined if migratory defaults proper to *Kif7* -/- cIN could moreover contribute to their abnormal cortical distribution.

### Kif7 invalidation affects the migratory behavior of cIN in co-cultures and organotypic slices

We thus compared the migratory behavior of *Kif7* -/- and WT cIN using an *in vitro* model previously established in the lab ^64^ to compare the dynamics of mutant and WT cIN migrating on a substrate of WT cortical cells (Fig.6A). WT and *Kif7* -/- E14.5 MGE explants from tdTomato expressing embryos were cultured on WT dissociated cortical cells (Fig. 6A1). Fluorescent migrating MGE cells were imaged using time-lapse video-microscopy and tracked over time (Fig.6A2). Dynamic parameters (speed during moves, stops, Fig. 6A3-A6) and trajectories (Fig. 6A7) were analyzed on movies. *Kif7* -/- cIN migrated slightly more rapidly than WT cIN (Fig. 6A3) mainly because the duration of their stops was significantly reduced (Fig. 6A4). The saltatory behavior of cIN was preserved (same frequency of fast nuclear movements, Fig. 6A5) despite a slight decrease of the speed of fast nuclear movements (Fig. 6A6). WT cIN migrated along pretty straight trajectories (Fig.6A2, left panel and Fig. 6A7, black curve) whereas *Kif7* -/- cIN showed reduced directionality persistence over time (Fig. 6A2, right panel and Fig. 6A7, blue curve). *Kif7* ablation thus affected the migratory behavior of cIN in a cell-autonomous manner and the migratory defaults of *Kif7 -/-* cIN evoked those of *Kif3a -/-* cIN whose primary cilium is non-functional (See Figure S6 in ^29^).

**Figure 6.**
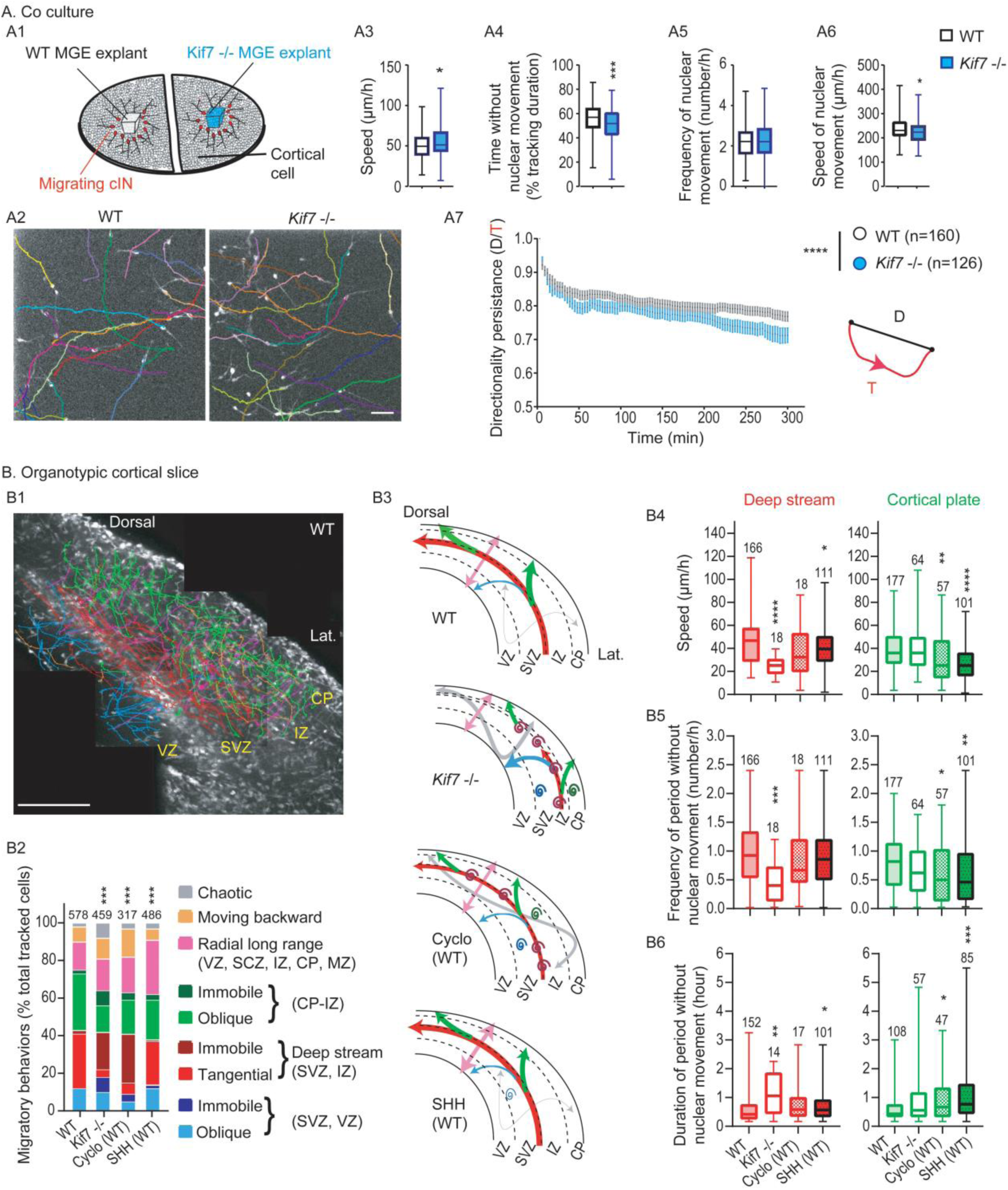
Dynamic behavior of migrating cIN in co culture and organotypic cortical slices. **(A)** To prepare co-cultures, MGE explants were dissected out of telencephalic vesicles at E14.5 from WT or *Kif7* -/- Nkx2.1-Cre;Rosa26-tdTomato embryos and placed on a substrate of WT E14.5 dissociated cortical cells (A1). After 24h in culture, MGE cells migrated centrifugally away from MGE explants on the substrate of cortical cells and were recorded for 7 hours.WT (n=160) and *Kif7* -/- (n=126) MGE cells were tracked manually using the MTrackJ plugin allowing to analyze migratory parameters (A2). Box and whisker plots indicate the speed (A3), the time without nuclear movement (A4), the frequency (A5) and speed of nuclear movements (A6). Statistical significance is assessed by Mann Whitney or Unpaired tests. ***, *P*<0.001 (A4); *, *P*=0.0308 (A3), *P*=0.0123 (A5). Directionality persistence was calculated as the ratio of the cell displacement (distance between the first and last positions of the cell) on the cell trajectory (A7, scheme on the right). Graph represented the mean directionality ratios at each time point over the first 5 hours of recording for the recorded cells (A7). Statistical significance is assessed using Two way ANOVA and reveals a significant effect of time and genotype (****, *P*<0.0001)]. Scale bar: 50 µm (A2). **(B)** tdTomato(+) cIN migrating in living cortical slices from Nkx2.1-Cre;Rosa26-tdTomato embryos were tracked manually using the MTrackJ plugin and their trajectories color-coded as shown in legend (B2) to characterize their preferred direction (tangential, oblique, radial, immobile) and cortical layer localization (VZ-SVZ, IZ, CP).The picture in B1 illustrates the z-projection of trajectories reconstructed in a control slice, superimposed to the last picture of the movie. Cortical interneurons migrating tangentially in the superficial migratory stream (MZ) could not be tracked because of high density. Graphs in B2 compare the percentage of each kind of trajectory recorded in the lateral cortex of control slices (control, see also Suppl. Movie S3), *Kif7* -/- slices (*Kif7* -/-, see also Suppl. Movie S4), control slices treated acutely with either murine SHH (SHH, see also Suppl. Movie S5) or cyclopamine (Cyclo, see also Suppl. Movie S6). Significance of the differences between the four distributions were assessed by a Chi-square test, X^2^ (24, *n*=1224), *P*=0.0004998, ***). All experimental conditions differed from the control (Fisher test, *P* =0.0004998, ***). Slices from three WT animals in control condition or treated with drugs and from three *Kif7* -/- animals were analyzed; the number of analyzed cells is indicated above bars. Schemes in B3 summarize the main results observed in each experimental condition. Trajectories are represented with the same color code as in B1, and line thickness is proportional to the percentage of cells exhibiting each type of trajectory. Immobile cells are figured by a coil. Box and whisker plots indicate the mean speed (B4), the frequency of period without nuclear movment (number/h) (B5) and the mean duration of period without nuclear movement (hour) (B6) for cIN migrating tangentially in the deep stream (red box and whisker plots, left) or to the cortical plate (green box and whisker plots, right). Statistical significance assessed by Krustkal-Wallis tests in each cluster. ****, *P*<0.0001; ***, B5 left *P*=0.0005; **, B4 right *P*=0.0088, B5 right, *P*=0.0037, B6 left *P*=0.0034; *, B4 left *P*=0.0233, B5 right *P*=0.0470, B6 left *P*=0.0134, B6 right *P*=0.0394. Number of analyzed cells is indicated on plots. VZ, ventricular zone; SVZ, subventricular zone; IZ, intermediate zone; CP, cortical plate; MZ, marginal zone. Scale bar: 300 µm.

We then analyzed the dynamic behavior of *Kif7* -/- cIN in the mutant cortex (Fig. 6B). In WT Nkx2.1-Cre;Rosa26-tdTomato transgenic slices, fluorescent cIN migrated tangentially from the PSB to the dorsal cortex in two main streams, the MZ and the lower IZ/SVZ (Fig. 6B1, Supplementary Movie S3). All along these pathways, cIN sporadically operated a tangential to oblique/radial migration switch to colonize the CP. High cell density in the MZ prevented cell monitoring and MZ cells were discarded from analyses. In the deep stream, a large proportion of imaged cells (59.3 %) either maintained tangentially oriented trajectories (28.7 %, red trajectories in Fig. 6B1,B3, left column of Fig. 6B2) or reoriented to the CP along oblique or radial trajectories (30.6 % green trajectories in Fig. 6B1,B3, left column in Fig. 6B2). Most of these cIN reached the CP surface and remained there. A small proportion of cIN moved from the deep stream to the ventricular side of the slice (11.7%, blue trajectories in Fig. 6B1,B3, left column in Fig. 6B2). A significant proportion (15.6 %) migrated radially over the cortex thickness, inverting their polarity in the MZ or at the ventricular surface (pink trajectories in Fig. 6B1,B3, left column in Fig. 6B2). Rare cIN in the VZ, deep stream and CP moved very short distances or did not move (3.6%, dark colors in the left column of Fig. 6B2). Finally, 8 % cIN (orange) moved backward to the ventral brain and 1.7 % (grey) showed no specific directionality or layer specificity and were classified as chaotic (Fig. 6B1,B3, left column in Fig. 6B2). The majority of cIN presented a characteristic saltatory behavior, alternating stops and fast moves ^64^. The dynamics of cells recorded in the deep stream and/or moving to the CP is shown in Fig. 6B4-6 (left column on histograms).

The trajectories followed by cIN in the lateral/latero-dorsal cortex of *Kif7 -/-* slices (where *CxCl12* mRNA was detected as shown in Fig. 5C) dramatically differed from those described above (compare WT and *Kif7* -/- columns in Fig. 6B2, schemes in Fig.6B3 and Supplementary Movies S3 and S4). The number of cIN migrating tangentially in the deep migratory stream and/or reorienting to the CP dropped drastically (17.8 % as compared to 59.3 % in WT slices) to the benefit of immobile cells (35.7 % as compared to 3.6 % in WT slices) especially in the SVZ/lower IZ. The proportions of cIN moving to the ventricle or to the CP were similar (10.2

% and 13.9 %, respectively, as compared to 11.7 % and 30.6 % in WT slices) suggesting that the radial asymmetry of cortical slices was lost for *Kif7 -/-* cIN. The same proportion of cIN migrated radially across the cortical layers as in WT slices (17.2 % versus 15.6 % in WT) but the proportion of cIN with chaotic trajectories strongly increased (7.8 % versus 1.7 % in WT slices). Another major change was a significant decrease of the migration speed (excluding immobile cIN) (Fig. 6B4) with less frequent but longer stops in *Kif7* -/- than in WT slices (Fig. 6B5,B6), confirming the dynamic parameters measured on co-cultures.

*Kif7* ablation has been shown to activate SHH transcriptional signaling by blocking the cleavage of GLI3 in GLI3-R repressor, or conversely to inhibit SHH transcriptional signaling by preventing the formation of GLI activators (GLI1/2)^3^. Our western blot analysis confirmed that the GLI3-R/GLI3-FL ratio dropped in the *Kif7 -/-* cortex according to the expected inhibition of the GLI3 processing, a result not observed in the MGE. Since *Kif7 -/*- cIN are born in the MGE and migrate in the cortex where SHH signaling is activated due to an abnormal GLI3 processing, we first examined whether the application of SHH on a WT forebrain slice could mimic the migratory defaults of *Kif7* -/- cIN in the mutant cortical slices. SHH application on control slices minimaly affected the trajectories of cIN (Fig. 6B2 compare WT and SHH (WT) columns). An increased proportion of cIN migrated radially across the cortical thickness (29 % compared to 15.6 % in WT slices) at the expense of cIN migrating from the deep stream to the CP (20.8 % instead of 30.6 % in WT slices). Frequent radial movements directed to the VZ (Fig. 6B2,B3, Supplementary Movie S5) did not evoke the migratory defaults of *Kif7 -/-* cIN in the mutant slices. Moreover, the frequency of chaotic trajectories did not increase (grey, Fig. 6B2) and the main dynamic alterations, a decreased speed (Fig.6B4) associated with an increased duration of stops (Fig. 6B6), were milder than those observed in *Kif7 -/-* cIN. On the contrary, the application of the SHH pathway inhibitor cyclopamine on WT cortical slices altered cIN trajectories in a way that resembled alterations in *Kif7-/-* slices (compare WT and cyclo (WT) columns in Fig. 6B2 and scheme in Fig. 6B3). For example, the proportion of immobile cIN and of cIN moving short distance reached 33.7 % as compared to 3.6 % in WT slices and 35.7% in *Kif7-/-* slices (Fig. 6B2). Cyclopamine application mainly affected the ability of cIN to move tangentially in the deep stream and to switch to the CP (Fig. 6B2,B3, Supplementary Movie S6). Nevertheless, the migration speed defects of *Kif7* -/- cIN and of cyclopamine treated cIN differed (Fig. 6B4).

In conclusion, *Kif7 -/-* cIN exhibited intrinsic migratory defaults resembling those of cIN with a primary cilium ablation and showed cortical trajectories defaults resembling those of cyclopamine treated WT cIN. *Kif7* ablation in cIN had the same effect than SHH signaling inhibition on cIN migratory behavior.

The strong influence of cyclopamine on the migration of cIN in WT cortical slices led us to determine the distribution of the endogenous activator SHH in the developing cortex.

### Distribution of Shh mRNA and SHH protein in the E14.5 cortex

*Shh* mRNA is strongly expressed in the mantle zone of the ganglionic eminences (MGE, CGE) and POA during the time cIN differenciate ^24,52^ but its expression level in cIN after they reach the cortex is controversial ^29,31,65^. We thus reexamined *Shh* mRNA expression in the forebrain and SHH protein distribution in cortical layers. *In* s*itu* hybridization (ISH) experiments performed at E13 confirmed the strong expression of *Shh* mRNA in the ventral forebrain, especially in the SVZ and mantle zone of the MGE (Fig. 7A1) in agreement with the SHH-dependent *Gli1* mRNA expression in a VZ subdomain of the MGE ^66^. In contrast, *Shh* transcripts were bearly detectable in the cortex. Using RNAscope, which has higher sensitivity and spatial resolution than standard ISH, we re-examined the expression pattern of *Shh* mRNA at E14.5. In the ventral forebrain, RNAscope *Shh* transcript detection showed similar expression pattern as ISH (Fig. 7A1,A3,B1,B2). In addition, *Shh* mRNA and *Lhx-6* mRNA, a cIN marker, were expressed in overlapping domains in the mantle zone of the MGE (Fig. 7A2-4). In the cortex, radially-oriented cells in the VZ/SVZ and tangentially oriented cells in the deepmigration stream expressed *Shh* mRNA at a very low level (one or two dots versus large cluster in the MGE) (Fig. 7C). Among the large number of cells expressing *Lhx-6* transcripts (Fig. 7C1,C2), only 5% co-expressed *Shh* mRNA (Fig. 7C3).

**Figure 7.**
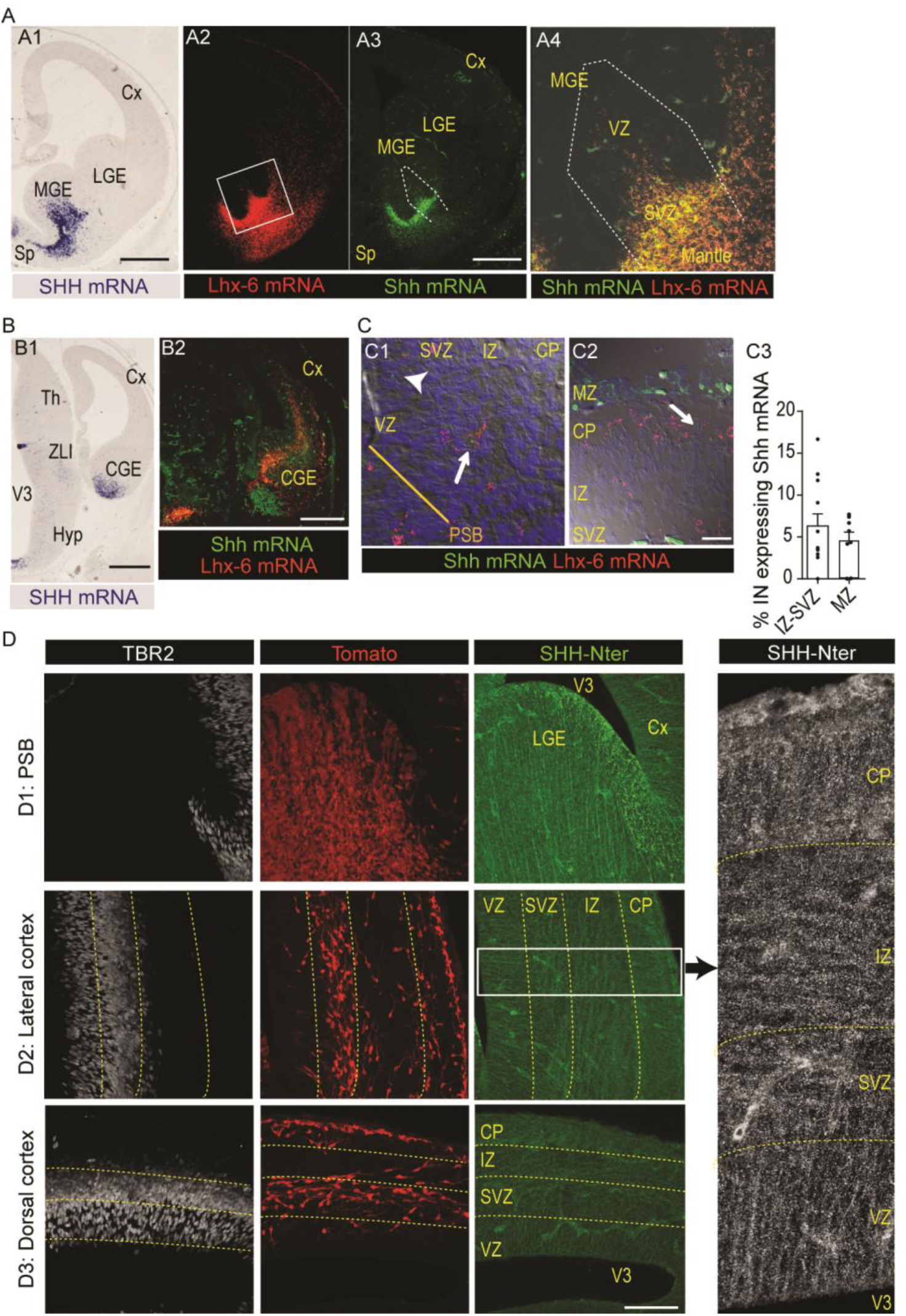
Expression of *Shh* transcripts (A-C) and distribution of SHH protein (D) in the E13.5-E14.5 forebrain. **(A,B)** Distribution in median (A) and caudal (B) coronal sections of Shh mRNA detected by *in situ* hybridization with an antisens *Shh* probe at E13.5 (A1,B1) and by RNAscope at E14.5 (A2-3,B2). *Shh* transcripts are strongly expressed in the medial ventral forebrain (SVZ and mantle zone of the MGE and septum, A1, A2), in the mantle zone of the CGE (B1,B2), in the zona limitans intrathalamica (ZLI in B1) and in the ventral midline of the 3^rd^ ventricle (V3 in B1). RNAscope further confirmed the strong expression of *Shh* mRNA (A3, green) in MGE and septum regions that strongly express *Lhx-6* mRNA (A2, red). Confocal observations in the SVZ and mantle zone of the MGE showed that *Shh* mRNA (green) is co-expressed with the *Lhx6* mRNA (red) in a significant number of cIN (yellow cells in A4). **(C)** Confocal analyses at higher magnification of the double detection by RNAscope of *Shh* and *Lhx-6* mRNA. Cells were identified on stacked images (Δz=1 µm) using Nomarski optic. In the lateral cortex close to the PSB (C1, z projection of 10 confocal planes) and in the dorsal cortex (C2, z projection of 10 confocal planes), a very small proportion of cells expressing *Lhx-6* mRNA also express *Shh* mRNA (white arrows in C1,C2). Counting in the deep stream (SVZ-IZ) and in the MZ is shown in graph C3 (9-17 fields in three sections). A few progenitors in the cortical VZ express *Shh* mRNA at very low level (arrowhead, C1). **(D)** SHH-Nter and TBR2 co-immunostaining of Nkx2.1-Cre/R26R-tdTomato brain coronal sections at E14.5. Representative confocal merged stacked images (Δz=0.2 µm; 48 images) in the pallium-subpallium boundary (PSB, D1), lateral cortex (D2) and dorsal (D3) cortex revealing SHH-Nter immunostaining in blood vessels and the presence of numerous bright dots all over the cortical neuropile. On the ventricular side of the PSB and of the lateral-most part of the LGE, SHH- Nter(+) brigth elements are aligned radially. In the lateral cortical neuropile, smaller bright dots align radially in the VZ, tangentially in the SVZ-IZ, and radially in the CP (see higher magnification on the right panel in which SHH-Nter immunostaining is shown in white and the contrast is increased). Cx, cortex; LGE, MGE and CGE, lateral, medial and caudal ganglionic eminence; CP, cortical plate; Hyp, hypothalamus; PSB, pallium-subpallium boundary; Sp, septum; Th, thalamus; V3, third ventricle; VZ, ventricular zone; SVZ, subventricular zone; IZ, intermediate zone; MZ, marginal zone. Scale bars: 500 µm (A, B), 20 µm (C), 100 µm (D).

Antibodies that recognize the N-ter part of SHH [SHH-Nter, e.g. “activated” SHH ^67^] strongly labeled in C57/Bl6 mouse brain the same regions known to strongly express SHH mRNA ^65,68,69^, such as the lateral wall of the third ventricle (Fig. S4A1) and the zona limitans intrathalamica at E12.5 (Fig. S4A2) and the choroid plexus and septum at E14.5 (Fig. S4B). SHH-Nter immunostaining was detected in the cortex at E12.5 and E14.5 (Fig. S4C,4D). To further investigate the expression of SHH in the cortical wall at E14.5, we performed immunostaining experiments in Nkx2.1-Cre/R26R-tdTomato embryos (Fig. 7D). SHH-Nter antibodies detected large bright dots that distributed radially away from the ventricle in the lateral LGE and at the ventricular angle (green dots in Fig. 7D1, upper panel). In the cortex, SHH-Nter antibodies immunostained bright tiny dots that show remarkable distribution (Fig. 7D2,D3). In the lateral (Fig. 7D2) and dorsal (Fig. 7D3) cortex, the bright SHH-ter immunopositive dots were aligned radially in the VZ and in the CP according to the radial arrangement of cortical progenitors and CP cells. In contrast, dots were aligned tangentially in the SVZ/IZ where cIN distribute among tangentially oriented growing axons. The intensity and density of positive dots appeared lower in the dorsal than in the lateral cortex suggesting a latero-dorsal gradient of distribution (compare Fig. 7D2 and 7D3). Some CP cells presented a diffuse cytoplasmic staining, which was not observed in the bottom layers. Together, our ISH and immunostaining experiments showed that the cortical SHH protein has probably an extrinsic origin as previously described at the early stage of cerebellum development ^70^.

## DISCUSSION

Our aim in the present paper was to better understand the developmental origin of functional abnormalities (e.g. epilepsy, cognitive deficits, …) in cortical circuits of human patients with *KIF7* mutations by characterizing the developmental abnormalities of the cerebral cortex in a *Kif7* -/- mouse model. We show that *Kif7* ablation affects the development of the two populations of cortical neurons, cortical plate neurons and GABAergic interneurons, in specific and distinct ways, accordingly with their dorsal and ventral origin in the telencephalon. The alterations of the CP development (lack of most subplate cells in the dorsal cortex, and delayed corticofugal projection) are remiscient of defaults previously described in GLI3 null mutants that lack most subplate neurons and pioneer cortical projections ^71^. The abnormal distribution of *Kif7* -/- cIN in the developing cortex partly reflects the structural and functional alterations of cortical layers on which cIN migrate (e.g. abnormal CXCL12 expression in the SVZ). However, we identify additional migration defaults of *Kif7* -/- cIN that recall those of cIN lacking a primary cilium or of wild type cIN treated with the SHH pathway inhibitor cyclopamine, suggesting that the SHH pathway activity is inhibited in *Kif7* -/- cIN. *Kif7* ablation thus alters differently the SHH pathway activity in CP and cIN subpopulations, in agreement with their dorsal or ventral origin.

### *Kif7* knock-out mice as a model to study the effect of *KIF7* mutations in human

KIF7 is a ciliary kinesin whose mutations in patients and ablation in mice alter the length rather than the structure of primary cilia. Primary cilium length was increased in patient fibroblasts ^10^, in MEF isolated from *Kif7* -/- and in the neural tube of *Kif7* -/- embryos at E10.5 ^73^. Owing that KIF7 regulates the trafficking and processing of GLI factors, the cilium-dependent SHH signaling was altered in both patient fibroblasts ^10^ and in *Kif7* -/- embryos ^3,73^ with GLI3-R decrease and GLI1/GLI2 up-regulation. These common features observed in cells from patients and in *Kif7* -/- mice models, as well as a previous study of the corpus callosum agenesis in *Kif7* mutant mice, which helped to dissect the mechanism at work in patients ^44^ allowed to propose that the *Kif7* -/- mouse is a good model to study the alterations that lead to clinical features in patients. Most *Kif7* -/- mice died at the end of gestation. However, the brain developed up to birth allowing to study its development.

All patients carrying mutations in the *KIF7* gene on both alleles have developmental delay/intellectual disability (ID) and some have seizures. Investigations on cerebral malformation by MRI have shown that some of them have alterations in the cerebral cortex including cortical atrophy ^9,12,13^ and pachygyria ^16^. Since detecting defects in the cerebral cortex is more demanding in MRI quality images than corpus callosum agenesis or molar tooth sign, we can speculate that the number of patients with cerebral cortex defects is underscored. Malformations of cortical development was recognized as causes of ID ^74^ and epilepsy ^75^ and the cellular defects underlying these pathologies could imply defects in cell proliferation, neuronal migration and differenciation that lead to improper excitatory/inhibitory balance, connectivity into the cortex or of the cortex with other brain structures. The mechanism of action of KIF7 to transduce SHH ciliary dependent signaling has been debated. It has been accepted that SHH activates KIF7 binding to microtubules and its accumulation to the tip of the cilium, making the kinesin motor domain the most essential domain for SHH transduction. A more recent study proposed that KIF7 is an immobile kinesin translocated to the cilium tip after SHH binding by a complex KIF3A/Kif3B/KAP ^76^ suggesting that domains in the KIF7 protein essential to its function are more complex. This could explain why any mutation, even point mutation in the Coiled-coil domain or in C-terminal part of the KIF7 protein induces clinical feature as severe as when mutation led to a highly truncated protein. Another explanation is that any mutation causes unproper folding of the protein probably inducing its degradation by the proteasome.

### KIF7 loss of function alters the cerebral cortex organization and the connectivity between the cortex and the thalamus

Our study demonstrates that in WT embryos, the levels of GLI3-FL and GLI3-R were much higher in the developing cortex than in the MGE and the cleavage of GLI3-FL in GLI3-R was more important in the cortex. Our observations are coherent with previous studies that identified a preponderant role of GLI3-R signaling to pattern the dorsal telencephalon and to regulate the proliferation and differenciation of neurons in the cortex ^41,42^. Moreover, the strong decrease of the cleavage of GLI3-FL in GLI3-R that we observed only in the cortex of *Kif7* -/- embryos suggests that the hyper SHH signaling resulting from this decreased cleavage affects the cortex, but not the MGE. The abnormal structure of the dorsal telencephalon in *Kif7 -/-* embryos (enlarged lateral ventricles, cortical wall folding and thinning) resembled the abnormal structure observed in ciliary mutants ^54,77^ and in GLI3 mutants ^78–80^. As in *Kif7-/-* embryos, a mild PBS disorganization and an abnormal differentiation of the caudal telencephalic vesicle and telo- diencephalic boundary that is dependent on SHH and GLI3 for its formation ^81^ were also reported in these mutants ^77,78^. The cellular disorganization observed in the developing cortex of *Kif7 -/-* embryos also recall defects of *Gli3* mutants, especially the impaired preplate and SP neurons differentiation, the abnormal cortical layer formation and abnormal axonal growth ^78–80^. Several of these traits have been observed in the cortex of the ciliary mutant Rfx3 -/- that exhibits also a decreased GLI3-R/GLI3-FL ratio in the dorsal telencephalon ^54^. Interestingly, the knock down of KIF7 in E14.5 cortical cells altered several developmental stages of cortical neurons, leading to a decrease of apical radial progenitors, basal IP divisions, and to impaired neuronal polarity and migration ^45^. Sporadic heterotopia as we observed in the cortex of *Kif7 -/-* embryos, have been reported in one patient ^16^. Cortical heterotopia is not a landmark of ciliopathy but has been also described at E11 in the forebrain of the *Ift88* hypomorphic mutant *cobblestone* (cbs, ^77^). And cellular alterations that should contribute to the formation of heterotopia, e.g. defective capacity to polarize and to migrate to the cortical plate, were described in post-mitotic neurons electroporated with shRNA libraries targeting ciliary proteins such as KIF7, BBS1, BBS10, NPHP8, ALMS1 ^45^. Heterotopias were also detected in a Joubert syndrom patient carrying a mutation in the *CELSR2* gene coding for a planar cell polarity protein ^82^ essential for neural progenitor cell fate decision, neural migration, axon guidance and neural maturation ^83,84^.

Beside delayed growth of cortical and thalamic axons, *Kif7* -/- embryos exhibited connectivity defects between the thalamus and cortex, with abnormal trajectories of axon bundles in the basal forebrain, at the telo-diencephalic junction and along the PSB. To our knowledge, the ectopic projection of cortical axons to the pre-optic area had not been observed in GLI3 mutants that exhibited instead an ectopic thalamic projection to the amygdala ^78,79^. In the ciliary mutants *Rfx3* -/-, ectopic cortical and thalamic axons were indeed observed in the basal telencephalon ^54^. Cortical neurons electroporated *in utero* with shRNAs to deplete KIF7 or ciliary proteins belonging to the BBSome, extended axons that showed abnormal fasculation and trajectories ^45^, demonstrating a cell-autonomous role of ciliary genes to control the axonal pathfinding. Finally, a large population of thalamic axons that had reached the cortex of *Kif7* -/- embryos showed a massive elimination at E18.5 in the caudal telencephalon. This massive loss paralleled the loss of CP cells, as previously observed in Rfx3 -/- and lnpp5e -/- mutants ^54^. Although disorganized thalamo-cortical connectivity had never been reported in patients with *KIF7* mutation or in patients with ciliopathy, thalamocortical connectivity is essential for cognitive performances in young infants ^85^ and its alteration leads to neurodevelopemental delay and later cognitive deficits ^86^. Moreover, a link beween ciliary function, cortex development and connectivity with the thalamus had been previously identified in mutant mice lacking the planar cell polarity proteins CELSR2 and FZD3. These mice display ciliary defects and an abnormal development of tracts between the thalamus and the cortex leading to an hypotrophy of the intermediate zone where thalamo-cortical and corticofugal axons navigate ^87,88^.

#### Extra-cortical origin of SHH

Beside its major function in forebrain patterning ^20,40^, SHH regulates the embryonic and perinatal cerebral cortex development ^27,28^. The origin of the morphogen in this brain structure remains obscure. In agreement with a recent set of data from Moreau et al. ^89^, we showed that cIN born in the MGE or POA no longer express SHH after they reach the developing cortex. In addition, the level of local *Shh* mRNA synthesis was extremely low in the cortex at E14.5. In contrast, SHH protein was detected in immunoposive dots distributed throughout the cortex. Dots aligned radially in the VZ and CP, and tangentially in the IZ. This cortical distribution is compatible with an extrinsic origin of SHH and a transventricular delivery as described in the cerebellar ventricular zone ^70^. SHH is indeed present in the cerebrospinal fluid and likely secreted by the choroid plexus, since *Shh* mRNA is expressed in the choroid plexus of mice from E12 to E14.5 ^90^. Accordingly, we showed here that the choroid plexus of the lateral ventricles was strongly immunopositive for SHH at E14.5. Interestingly, the CDO/BOC co-receptor - that favors SHH transport - was also detected along radially oriented structures by immunostaining in the mouse embryonic cortex (not illustrated). SHH could thus be transported away from the ventricular zone and/or meninges in vesicular structures able to diffuse accross the cortical thickness. Accordingly, it has been shown that SHH transport needs ESCRT-III member protein CHMP1A ^15^ and SHH related proteins are found in exosomes secreted from Caenorhabditis elegans epiderman cells ^72^, Drosophila wing imaginal discs ^88^, chick notochord cells and human cell lines ^91^. How SHH is transported throughout the cortex should be further explored. It has already be shown that SHH is present in multivesicular bodies, in neurites and filipodia in the postnatal hippocampus and cerebellum, and in vesicles located inside and outside of hippocampal neurons in culture ^92^.

#### *Kif7* deletion leads to defects in cortical interneuron migration

The defects induced by gene mutations responsible for primary cilium dysfunction, including *Kif7* deletion, have been examined in principal neurons born in the dorsal telencephalon, which organize the cortical layers ^93,94^. However, the cIN are born ventrally in the MGE where the expression levels of GLI3-R and GLI3-FL are extremely low in both WT and *Kif 7-/-* embryos. cIN leave the MGE and then migrate to the cortex during the embryonic development. Their primary cilium is functional and can transduce local SHH signals to influence their directionality, especially the tangential to radial reorientation of their trajectories toward the cortical plate ^29^. In the present study, we showed that the abnormal distribution of cIN in the mutant cortex of *Kif7* -/- embryos resulted from cell autonomous defects and from extrinsic cortical defects, especially the transient lack of CXCL12 expression in the dorsal cortex at E14.5. Interestingly, *Kif7* ablation affected the migratory behavior of cIN cultured on a WT cortical substrate in a similar way as did the ablation of *Kif3a* ^29^, another ciliary gene required for the transduction of SHH signals ^95^. Accordingly, cyclopamine application on WT cortical slices altered the migratory behavior of cIN in a way that mimicked the migratory behavior of Kif7 -/- cIN. We thus concluded from these observations that *Kif7* ablation inhibited the Shh pathway activity in cIN. By consequence, *Kif7* -/- cIN, which normally differentiate in a ventral telencephalic domain expressing high SHH level, present dynamic and trajectories alterations that mimics those of cIN lacking SHH signalling. Therefore *Kif7* ablation in MGE-derived cIN does not activate SHH signalling by decreasing the GLI3-R/GLI3-FL ratio, as observed in the dorsal telencephalon. On the contrary, Kif7 ablation in MGE-derived cIN likely inhibits SHH signalling by a mechanism independent of GLI3 processing that was unchanged in the MGE of *Kif7 -/-*embryos (Fig.2). In addition to this cell autonomous defect that impaired cIN migration, we show that the lack of CXCL12 expression in the dorsal cortex transiently and locally modified the directionality of Kif7-/- cIN altering thereby the dorsal cortex colonization by cIN in *Kif7* mutants ^59,96^.

Interestingly, acute SHH application on WT cortical slices slightly increased the percentage of radially migrating cIN. In contrast, cyclopamine that inhibited the SHH pathway blocked the migration of a large population of cIN in the deep cortical layers demonstrating that endogeneous cortical SHH is likely to activate the SHH pathway in migrating cIN to control their trajectories.

In conclusion, using the *Kif7* -/- murine model, we show that the *Kif7* deletion is responsible for a large range of developmental defects in the cerebral cortex, which include: i) alteration of the inhibitory interneurons migration by a combination of intrinsic and extrinsic factors, ii) abnormal formation of cortical layers by the cortical plate neurons and iii) major pathfinding defects in axonal projections between the cortex and the thalamus. Patients carrying mutations in the *KIF7* gene are classified as ciliopathic patients and display developmental delay, ID and epilepsy. These clinical features could thus be caused by an abnormal distribution of excitatory and inhibitory cortical neurons leading to impaired excitatory/inhibitory balance, and by the development of abnormal connections in the cerebral cortex and with others brain structures. By demonstrating focal heterotopia and abnormal thalamo-cortical connectivity in the murine model, we hope to open a new field of clinical investigations using MRI and tractography to identify such defects in patients carrying mutations in the *KIF7* gene.

## MATERIAL AND METHODS

### Mice

E14.5 embryos of the swiss (for *in situ* hybridization and RNAscope) and C57/Bl6J (for sonic hedgehog immunostaining) strains were obtained from pregnant females purchased at JanvierLabs (France). Male and female wild type (WT) and *Kif7* -/- animals (E12.5, E14.5, E16.5, E18.5 and P0) expressing tdTomato or not in the MGE-derived cIN (Nkx2.1- Cre;Rosa26-tdTomato) were generated in our animal facility by mating male *Kif7* +/-; Nkx2.1- Cre;Rosa26-tdTomato and female *Kif7* +/- mice to produce embryos and P0. Thirty four pregnant females were used. *Kif7* +/- mice are a generous gift of Pr Chi-Chung Hui (the Provider, SickKids Hospital, Toronto, USA) and Prof Bénédicte Durand (the Transferor, CNRS, Lyon, France). Mice were genotyped as described previously ^3^. The day of the vaginal plug was noted E0.5. Experiments have been validated and approved by the Ethical committee Charles Darwin (C2EA-05, authorized project 02241.02) and mice were housed and mated according to European guidelines. Both male and female animals were used in this study. Sex is not considered as a biological variable in embryos.

### Brain lysate preparation and immunoblotting

Cortex and MGE from E14.5 WT and *Kif7* -/- embryos were dissected and frozen at -80°C until use. Samples were sonicated in 80µl Laemmli buffer and heated for 10 min at 37°C after adding 2.5% β-mercapto ethanol. Proteins were loaded on SDS-PAGE 7% gels (BIO-RAD) and transferred to 0.45 μm Nitrocellulose membranes (BIO-RAD). Membranes were cut to detect signals from proteins above and below 55 KDa and were blocked with 50 g/l non-fat dry milk in PBS/0.1% Tween 20 for 1h at RT, incubated with primary antibodies in the same solution O/N at 4°C, 1 h at RT with appropriate IRDye-conjugated secondary antibodies, and imaged and quantified using ChemiDocMP Imaging System (BIO-RAD). Commercial antibodies anti GLI3 (R&D Sytem-Biotechne, goat, #AF369, 1:200), anti-actin (Millipore, mice, #MAB1501, 1:10.000), Donkey-anti-goat-IR800 (Advansta, R-05781, 1:20.000), Goat-anti-mouse-StarBight Blue 700 (BIO-RA, 12004159, 1:2.500). Band intensities were quantified using gel analysis of Fiji software.

### Immunohistochemistry and imaging

Heads of embryos and P0 animals were fixed by immersion in 0.1% picric acid /4% paraformaldehyde (PAF) in phosphate buffer (PB) for 4 hours and then in 4% PAF in PB overnight. After PBS washes, brains were dissected, included in 3% type VII agarose (Sigma, A0701) and sectionned at 70 μm with a vibratome. Immunostaining was performed on free-floating sections as explain previously ^29^ . Primary antibodies were goat anti SHH-Nter (1:100, R&D system AF464), goat anti Netrin G1a (NG1a) (1:100, R&D system AF1166), rabbit anti TBR1 (1:1000, Abcam ab31940), rabbit anti TBR2 (1:1000, Abcam ab23345), rabbit anti PAX6 (1:100, clone poly19013, Covance PRB-278P), rabbit anti GSH2 (1:2000, Millipore ABN162), chicken anti MAP2 (1:500, Novus, NB30213), mouse anti SMI-32 (1:500, Covance SMI-32R), and rat CTIP2 (1:1000, Abcam ab18465). Primary antibodies were incubated in PGT (PBS/gelatine 2g per L/TritonX-100 0.25%) with 0.1% glycine and 0.1% lysine for anti Netrin G1a antibodies and 3% BSA for anti SHH-Nter antibodies. Primary antibodies were revealed by immunofluorescence with the appropriate Alexa dye (Molecular Probes) or Cy3 (Jackson laboratories) conjugated secondary antibodies diluted in PGT (1:400). Bisbenzimide (1/5000 in PBT, Sigma) was used for nuclear counterstaining. Sections were mounted in mowiol/DABCO (25mg/mL) and imaged on a macroscope (MVX10 olympus) or an epifluorescence microscope (LEICA DM6000) using X40 and X63 immersion objectives, or a confocal microscope (Leica TCS SP5) using a x40 objective. Double immunostaining were performed when possible and images were representative of what was observed in more than 4 animals per genotype except for heterotopia. Macroscop images were treated to remove background around sections using wand tool in Adobe Photoshop 7.0 software. On epifluorescence and confocal microscop images, acquisition parameters and levels of intensity in Adobe Photoshop 7.0 software were similar for WT and *Kif7* -/-. Merged images were acquired on the same section except for TBR1 with TBR2 acquired on adjacent sections. The fluorescence intensity along the depth of the cortex was assessed using the plot profile function of ImageJ under a 500 pixels wide line starting in the ventricle to the surface of the brain.

### In situ hybridization

Specific antisense RNA probe for *Shh*, *Gli1* and *CxCl12* genes (gift of Marie-Catherine Tiveron) were used for in situ hybridization analyses. DIG-probes were synthesized with a labeling kit according to the manufacturer’s instructions (Roche, France). *In situ* hybridization was performed according to Tiveron et al ^97^ with few modifications (see Supplementary material). Sections were mounted on glass slides, dried, dehydrated in graded ethanol solutions, cleared in xylene and coverslipped with Eukitt.

### RNAscope

Embryonic E14.5 heads were cuted in cold Leibovitz medium (Invitrogen) and immersion fixed in cold 4% (w/v) paraformaldehyde (PFA) in 0.12 m phosphate buffer, pH 7.4, overnight. Brains were then cryoprotected in PFA 4% /sucrose 10 %, embedded in gelatin 7.5% /sucrose 10% at 4°C and frozen. Biological samples were kept at −80°C until coronally sectioned at 20 μm with cryostat. RNAscope experiment was performed according manufacturer’s instructions (RNAScope^®^ Multiplex Fluorescent V2 Assay, ACDbio) after a pre-treatment to remove gelatin. Two different probes/channels were used (C1 for Shh, C2 for Lhx-6). Probes were diluted according manufacturer’s instructions (1 Vol of C2 for 50 vol of C1) and dilution of the fluorophores (Opal 520 and 570) was 1:1500. Sections were mounted in mowiol/DABCO (25mg/mL) and imaged on a confocal microscop (Leica TCS SP5) using an immersion X10 or X63 objectives on 10 µm stacks of images (1 image /µm).

### DiI experiments

Tiny crystals of DiI (1,1′-dioctadecyl-3,3,3′3′-tetramethylindocarbocyanine perchlorate) were inserted in the dorsal or lateral cortex of WT and *Kif7* -/- embryonic brains fixed at E14.5 by immersion in 4% PAF. Injected brains were stored in 4% PAF at room temperature in the dark for several weeks to allow DiI diffusion. Before sectioning, brains were included in 3% agar. Coronal sections 60 μm thick were prepared with a vibratome and collected individually in 4% PAF. Sections were imaged with a macroscope (MVX10 Olympus).

### Whole brain clearing and imaging

E14 and E16 embryos (WT, n=5; *Kif7* -/-, n=6) were collected in cold L15 medium, perfused transcardially with 4% PAF using a binocular microscope. Brains were dissected and postfixed 3 days in 4% PAF at 4°C, and stored in PB at 4°C until clearing. All buffer solutions were supplemented with 0.01% Sodium Azide (Sigma-Aldrich). Whole brain immunostaining and clearing was performed at RT under gentle shaking according to a modified version of the original protocol ^98^. Briefly, perfused brains were dehydrated in graded methanol solutions (Fischer Scientific) (20, 50, 80% in ddH_2_O, 2 × 100%, 1h30 each), bleached overnight in 3% hydrogen peroxide (Sigma-Aldrich) in 100% methanol and rehydrated progressively. After PBS washes, brains were blocked and permeabilized 2 days in PBGT (0.2% gelatin (Merck), 0.5% triton X-100 (Sigma-Aldrich) in PBS). Brains were incubated 3 days at 37°C in primary goat anti Netrin G1a (1:100, R&D system AF1166) and rabbit anti TBR1 (1:1000, Abcam ab31940) antibodies. After 1 day of PBGT washes, brains were incubated 1 day at 37°C in secondary 647 donkey anti-goat (1:1000, Molecular probes,) and cy3 donkey anti-rabbit (1:1000, Jackson laboratories) antibodies. Immunolabeled brains were washed 1 day in PBS, embedded into 1.5% low-melting agarose (type VII, Sigma) in 1% ultra-pure Tris-acetate-EDTA solution, placed in 5-ml tubes (Eppendorf, 0030119452) and dehydrated 1h in each methanol baths (50%, 80% and 2 × 100%). Samples were then incubated for 3h in 33% methanol / 66% dichloromethane (DCM, Sigma-Aldrich), washed in 100% DCM (2×30 min) and incubated overnight (without shaking) in dibenzyl ether (DBE). Brains were stored in DBE at RT. Cleared samples were imaged on a light-sheet microscope (LaVision Biotec, x6.3 zoom magnification) equipped with a sCMOS camera (Andor Neo) and Imspector Microscope controller software. Imaris (Bitplane) was used for 3D reconstructions, snapshots and movies.

### Brain slices

Brains of E14.5 WT and *Kif7* -/- transgenic Nkx2.1-cre, tdTomato embryos were dissected in cold L15 medium, embedded in 3% type VII agarose and sectioned coronally with a manual slicer as explained in ^29^. Forebrain slices 250 µm thick were transferred in Millicell chambers and cultured for 4 hours in F12/DMEM medium supplemented with CFS 10% in a CO2 incubator (Merck Millipore) prior to pharmacological treatment and recording. Slices were incubated in drug for 2 hours before transfer in culture boxes equipped with a bottom glass coverslip for time-lapse imaging.

### Pharmacological treatment

Either recombinant Mouse SHH (C25II), N-terminus (464-SH- 025, R&D systems) at 0.5µg/mL final or cyclopamine (C4116, Sigma) at 2.5 µM final were added to the culture medium of slices and renewed after 12 hours. Recombinant SHH was reconstituted at 100 µg/mL in sterile PBS with BSA 0.1%. The stock solution of cyclopamine was 10 mM in DMSO and the culture medium of control experiments contained 1/4000 DMSO (vehicle). Control experiments in culture medium with and without vehicle did not differ and were analyzed together.

#### Co-cultures

Co-cultures were performed on polylysine/laminin-coated glass coverslips fixed to the bottom of perforated Petri dishes in order to image migrating MGE cells. Brains were collected in cold PBS at embryonic day E14.5. Cortices and MGE explants were then dissected in cold Leibovitz medium (Invitrogen). Cortices were mechanically dissociated. Dissociated cortical cells were cultured on the coated glass coverslips and left in the incubator (37 °C, 5% CO2) for 1 hours. A wound divided the substrate in two-halves on which WT and *Kif7* -/- MGE explants from littermate embryos were placed (Fig. 6A). Co-cultures were cultured for 24h and imaged.

### Videomicroscopy and image processing

Slices and co-cultures were imaged on an inverted microscope (Leica DMI4000) equipped with a spinning disk (Roper Scientific, USA) and a temperature-controlled chamber. Multi-position acquisition was performed with a Coolsnap HQ camera (Roper Scientific, USA) to allow the recording of the whole cortex. Images were acquired with a X20 objective (LX20, Fluotar, Leica, Germany) and a 561 nm laser (MAG Biosystems, Arizona). Z-stacks of 30 μm were acquired 50 µm away from the slice surface, with a step size of 2 μm and a time interval of 2 or 5 minutes for at least 21 hours. Acquisitions were controlled using the Metamorph software (Roper Scientific, USA). Cell trajectories were reconstructed on movies by tracking manually the cell rear with MTrakJ (ImageJ plugin, NIH, USA) and were clustered according to their position in cortical layers and to their orientation (see details in Fig. 6). The migration speed of cells, the frequency and duration of pauses were extracted from tracking data using excel macros. The directionality of cells was analysed along time using a macro from ^99^.

### Study design

WT group was compared to *Kif7* -/- group and to pharmacological treatment groups for brain slice experiments. The experimental unit was a single animal for immunohistochemistry analysis and individual cell for videomicroscopy analysis.

### Statistical analysis

All data were obtained from at least three independent experiments and are presented as mean ± SEM (standard error of mean). Statistical analyses were performed with the GraphPad Prism software or R. Statistical significance of the data was evaluated using the unpaired two-tailed *t* test, the Mann–Whitney test, the Chi2 test or the Two-way ANOVA test followed by a post-hoc test. Data distribution was tested for normality using the D’Agostino and Pearson omnibus normality test. Values of *P*<0.05 were considered significant. In figures, levels of significance were expressed by * for *P*<0.05, ** for *P*<0.01, *** for *P*<0.001 and **** for *P*<0.0001.

## Data availabitily

The data presented in this study are available from the corresponding author upon request without undue reservation.

## Acknowledgements

Marie-Christine Tiveron is acknowledged for the gift of the CXCL12 expression vector. We thank all members of the Métin’s Team for constructive discussions. We gratefully acknowledge the Imaging plateform facility of the Fer à Moulin Institute for the use of their microscopes, and the animal facility of the Fer à Moulin Institute for animal care and breeding.

## Author contributions

CM and JM conceptualized the project and supervised the work. MP, VG, SSL, JP, AM, NR, SL, CM and JM perfomed experiments. MP, SSL, AM, NR, CM and JM analyzed the data. MP, VG, SSL, CM and JM wrote the paper.

**We declare** no Competing interest.

## SUPPLEMENTARY MATERIAL

### MATERIAL AND METHODS

#### In situ hybridization

Fixed embryos were cryoprotected overnight in PBS with 10% sucrose, embedded in OCT (Tissue-Tek, Miles, Elkhart, IN) and frozen on dry ice. 10 µm cryostat sections were thaw-mounted on Superfrost slides (Menzel-Gläser), left to dry at room temperature (RT), and stored at -80°C. Thawed sections were treated two X 10 min in RIPA buffer (150 mM NaCl, 1% NP-40, 0.5% Na deoxycholate, 0.1% SDS, 1 mM EDTA, 50 mM Tris, pH 8.0), postfixed in 4% PFA for 10 min at RT, and washed three X 5 min with PBS. The slides were then transferred in 100 mM triethanolamine, pH 8.0 for 2 min, and then acetylated for 10 min at RT by adding dropwise acetic anhydride (0.25% final concentration) while being rocked, and washed again three X 5 min in PBS. The slides were prehybridized briefly with 500 µl of hybridization solution (50% formamide, 5x SSC, 5x Denhardt’s, 500 mg/ml herring sperm DNA, 250 mg/ml yeast RNA) and hybridized overnight at 70°C with the same solution in the presence of the heat-denatured DIG-labeled RNA probe. The following day, slides were washed in posthybridization solution (50% formamide, 2x SSC, 0.1% tween20) at 70°C first until coverslips slid off, then twice for 60 min at 70°C and finally at RT for 5 min. Slides were washed with buffer 1 (100 mM maleic acid, pH 7.5, 150 mM NaCl, 0.05% Tween 20), blocked for 30 min in buffer 2 (10% heat-inactivated horse serum in buffer 1), incubate overnight at 4°C with alkaline phosphatase-coupled anti-DIG antibody (Roche Diagnostics, Mannheim, Germany) diluted 1:1000 in buffer 2, rinsed twice for 5 min with buffer 1, and equilibrated for 30 min in buffer 3 (100 mM Tris, pH 9.5, 100 mM NaCl, 50 mM MgCl2). The signal was visualized by a color reaction using 250 µl of buffer 4 per slice [6.6 µl/ml NBT (4-nitroblue tetrazolium chloride, Roche Diagnostics), 3.3 µl/ml BCIP (5-bromo-4-chloro-3-indoyl- phosphate, Roche Diagnostics) in buffer 3]. The color reaction was allowed to develop in the dark at RT during few hours and was stopped with PBS.

**Figure S1.**
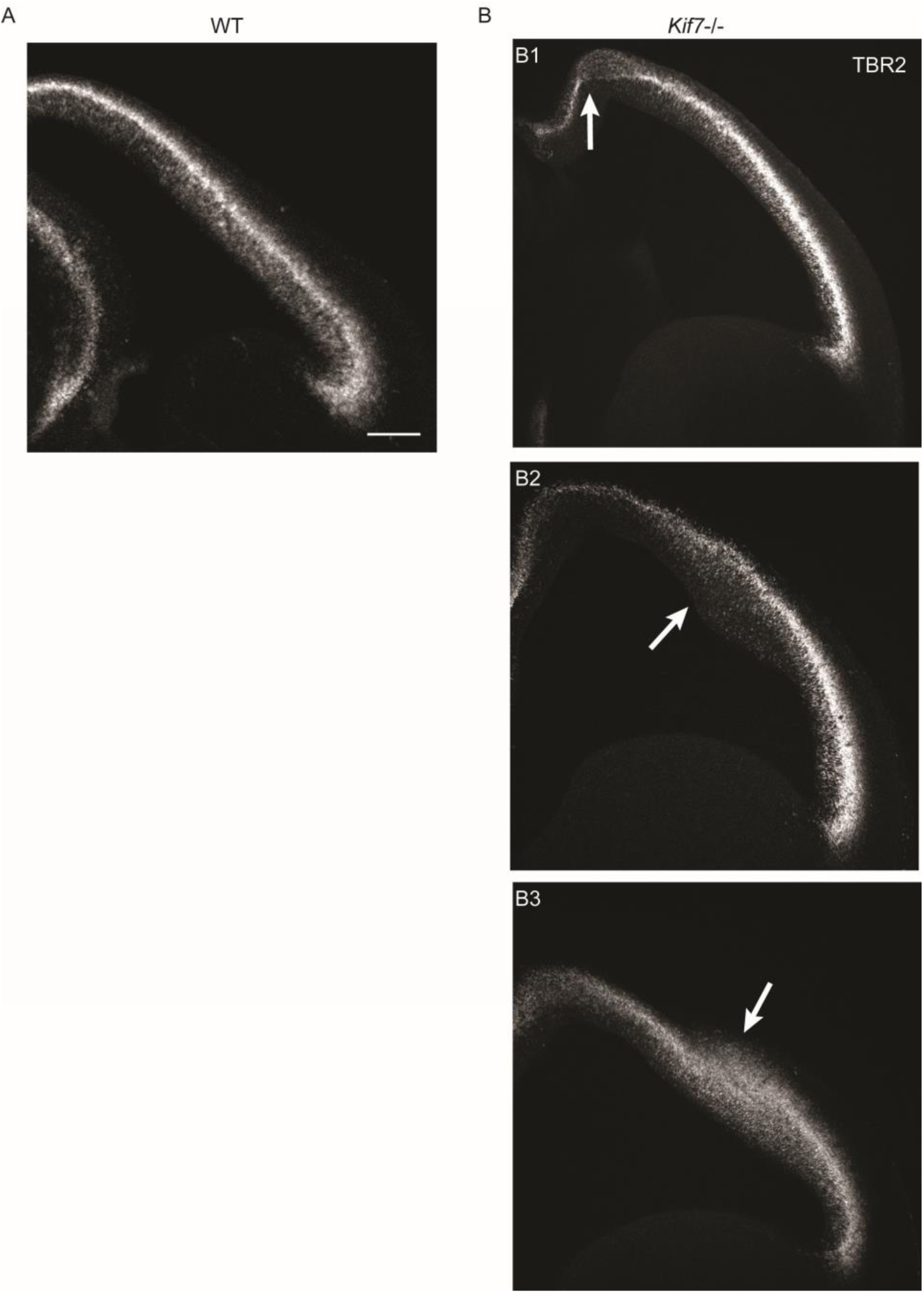
*Kif7* deletion is associated with cortical heterotopia at E14.5. **(A)** On coronal sections of the telencephalon of WT animals, TBR2(+) cells form a well-defined layer restricted to the cerebral cortex. Most TBR2(+) cells are densely packed in the SVZ, whereas some TBR2(+) cells are dispersed in the ventricular zone. **(B)** In about 20% of *Kif7* -/- embryos, heterotopia identified by a disorganization of the TBR2(+) layer were observed in various regions of the cerebral cortex, either dorsal (B1) or lateral (B2,B3). Depending on heterotopia, TBR2(+) were either displaced toward the brain surface (B2) or toward the ventricle (B3). Scale bar: 200 µm.

**Figure S2.**
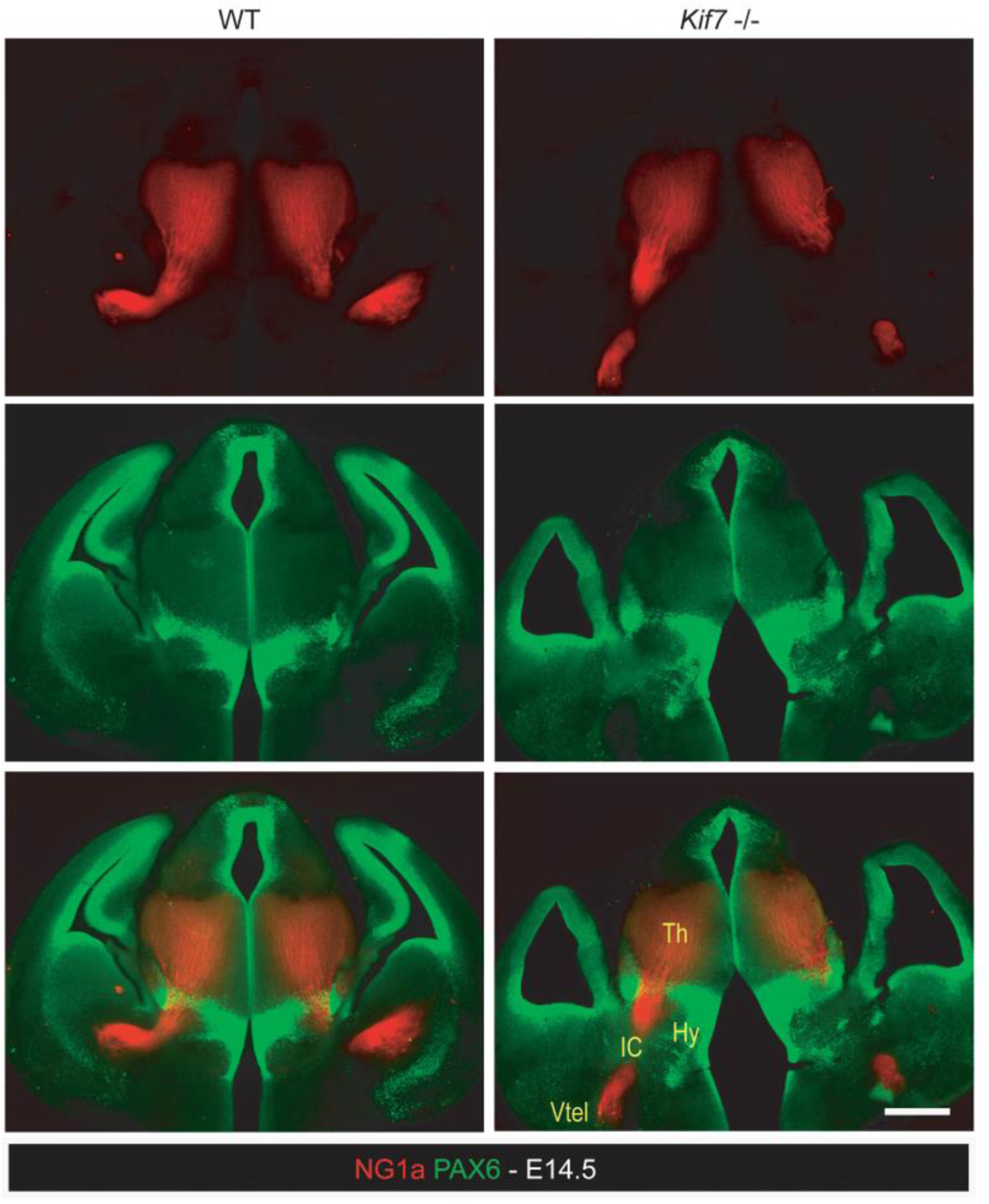
Alterations of the thalamo-cortical projection at E14.5 in *Kif7* -/- brains. Immunostaining of coronal brain sections at a caudal level with NG1a (red) and PAX6 (green) antibodies shows that TCA mistargeting in the ventral telencephalon is not related to abnormal PAX6 expression in the zona incerta between the dorsal and ventral thalamus. Scale bar: 200 µm.

**Figure S3.**
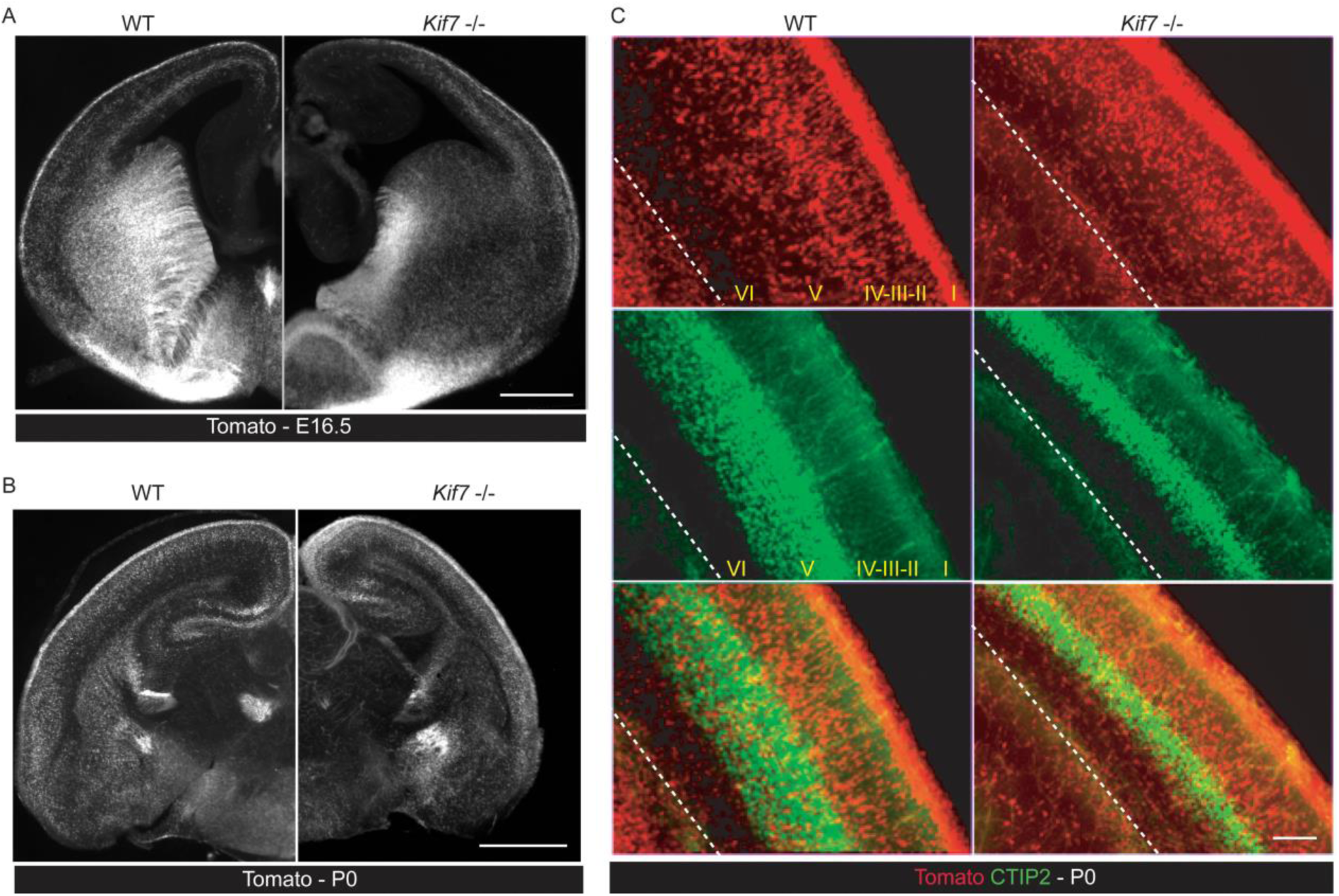
cIN migration up to birth. **(A)** At E16.5, the delay observed in the ability of cIN to migrate toward the dorsal cortex onserved at E14.5 persisted, however, some cIN were able to migrate forward but no longer following the WT pattern of tangential migration. **(B,C)** At birth (P0), cIN had colonized to dorsal cortex in Kif7 -/- embryos (C) but are abnormally distributed in the *Kif7* -/- cortex: their density is strongly incresed in the supragranular cortical layers that do not express CTIP(+) principal neurons (green) (C). Scale bar: 500 µm (A,B), 100 µm (C).

**Figure S4.**
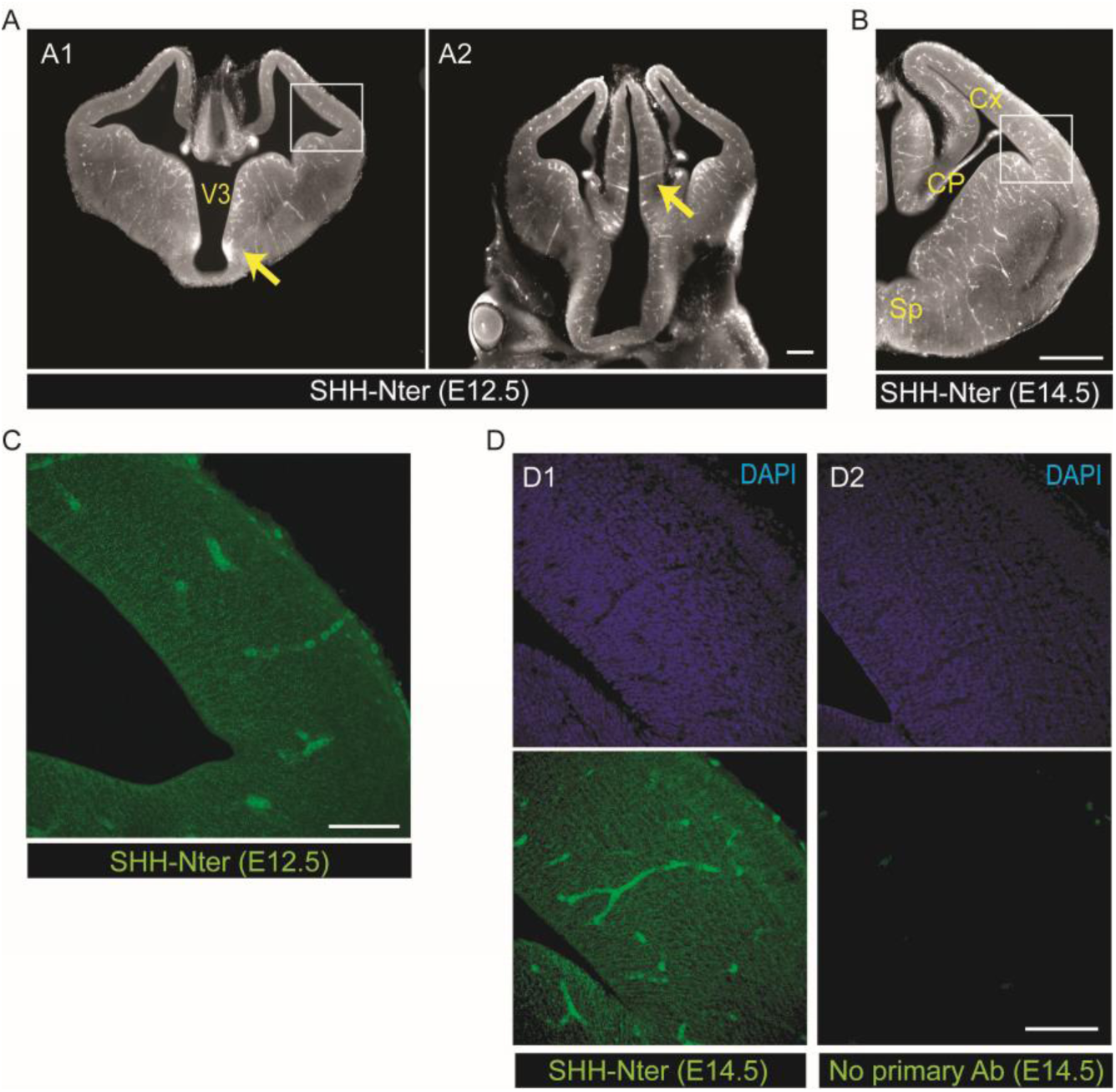
SHH-Nter immunostaining specifity. **(A,B)** Representative pictures of SHH-Nter immunostaining of C57B6 mice brain coronal sections at E12.5 (A) and E14.5 (B) imaged with a macroscope. High signal is observed along the third ventricle (A1, arrow) and in the ZLI (A2, arrow) at E12.5 and in the choroid plexus and the septum at E14.5 (B) whereas a faint signal is observed in the cortex. **(C)** Representative picture of SHH-Nter immunostaining in the cortex at E12.5. Z stacks of 48 images confocal images (Δz=0.2 µm) reveals the presence of numerous bright dots in the cortical neuropile with a gradient of density from the ventricular surface to the surface of the brain. **(D)** Coronal sections of E14.5 brain are labeled with SHH-Nter antibodies (D1) or only with the secondary antibodies (D2). Confocal Z stacks of 48 images (Δz=0.2 µm) reveal a punctiform signal all over the cortical neuropile and in some cells in the cortical plate (D1) whereas no signal is observed in sections immunolabeled only with the secondary fluorescent antibody (D2). CP, choroid plexus; Cx, cortex; Sp, septum; V3, third ventricle; ZLI, zona intra-thalamica. Scale bar: 250 µm (A,B), 100 µm (C,D).

**Movie S1 (related to Fig. 3).** 3D maximum intensity projection of thalamo-cortical axons in E14.5 WT brain. Whole-mount iDISCO+ immunolabeling for NG1a (in red) and TBR1(+) neurons (in green) imaged with light sheet microscopy.

**Movie S2 (related to Fig. 3).** 3D maximum intensity projection of thalamo-cortical axons in E14.5 *Kif7* -/- brain. Whole-mount iDISCO+ immunolabeling for NG1a (in red) and TBR1(+) neurons (in green) imaged with light sheet microscopy.

**Movie S3 (related to Fig. 5).** Migration of tdTomato expressing WT MGE cells. Forebrain slices 250 µm thick are imaged on an inverted microscope. Time-lapse between frames is 10 minutes.

**Movie S4 (related to Fig. 5).** Migration of tdTomato expressing *Kif7* -/- MGE cells. Forebrain slices 250 µm thick are imaged on an inverted microscope. Time-lapse between frames is 10 minutes.

**Movie S5 (related to Fig.5).** Migration of tdTomato expressing WT MGE cells treated with mouse SHH. Forebrain slices 250 µm thick are imaged on an inverted microscope. Time-lapse between frames is 10 minutes.

**Movie S6 (related to Fig.5).** Migration of tdTomato expressing WT MGE cells treated with cyclopamine. Forebrain slices 250 µm thick are imaged on an inverted microscope. Time-lapse between frames is 10 minutes.

## References

1. Park SM, Jang HJ, Lee JH. Roles of Primary Cilia in the Developing Brain. Front Cell Neurosci. 2019;13:218. doi:10.3389/fncel.2019.00218

2. Haycraft CJ, Banizs B, Aydin-Son Y, Zhang Q, Michaud EJ, Yoder BK. Gli2 and Gli3 Localize to Cilia and Require the Intraflagellar Transport Protein Polaris for Processing and Function. Barsh G, ed. PLoS Genet. 2005;1(4):e53. doi:10.1371/journal.pgen.0010053

3. Cheung HO, Zhang X, Ribeiro A, et al. The kinesin protein Kif7 is a critical regulator of Gli transcription factors in mammalian hedgehog signaling. Science signaling. 2009;2(76):ra29. doi:10.1126/scisignal.2000405

4. Endoh-Yamagami S, Evangelista M, Wilson D, et al. The mammalian Cos2 homolog Kif7 plays an essential role in modulating Hh signal transduction during development. Current biology : CB. 2009;19(15):1320–1326. doi:10.1016/j.cub.2009.06.046

5. Liem KF, He M, Ocbina PJR, Anderson KV. Mouse Kif7/Costal2 is a cilia-associated protein that regulates Sonic hedgehog signaling. Proc Natl Acad Sci USA. 2009;106(32):13377–13382. doi:10.1073/pnas.0906944106

6. Ingham PW, McMahon AP. Hedgehog Signalling: Kif7 Is Not That Fishy After All. Current Biology. 2009;19(17):R729–R731. doi:10.1016/j.cub.2009.07.060

7. Pedersen LB, Akhmanova A. Kif7 keeps cilia tips in shape. Nature cell biology. 2014;16(7):623–625. doi:10.1038/ncb2997

8. Han Y, Wang B, Cho YS, et al. Phosphorylation of Ci/Gli by Fused Family Kinases Promotes Hedgehog Signaling. Developmental Cell. 2019;50(5):610–626.e4. doi:10.1016/j.devcel.2019.06.008

9. al-Gazali LI, Bakalinova D. Autosomal recessive syndrome of macrocephaly, multiple epiphyseal dysplasia and distinctive facial appearance. Clinical dysmorphology. 1998;7(3):177–184. doi:10.1097/00019605-199807000-00004

10. Putoux A, Thomas S, Coene KL, et al. KIF7 mutations cause fetal hydrolethalus and acrocallosal syndromes. Nature genetics. 2011;43(6):601–606. doi:10.1038/ng.826

11. Putoux A, Nampoothiri S, Laurent N, et al. Novel KIF7 mutations extend the phenotypic spectrum of acrocallosal syndrome. Journal of medical genetics. 2012;49(11):713–720. doi:10.1136/jmedgenet-2012-101016

12. Ali BR, Silhavy JL, Akawi NA, Gleeson JG, Al-Gazali L. A mutation in KIF7 is responsible for the autosomal recessive syndrome of macrocephaly, multiple epiphyseal dysplasia and distinctive facial appearance. Orphanet journal of rare diseases. 2012;7:27. doi:10.1186/1750-1172-7-27

13. Walsh DM, Shalev SA, Simpson MA, et al. Acrocallosal syndrome: identification of a novel KIF7 mutation and evidence for oligogenic inheritance. European journal of medical genetics. 2013;56(1):39–42. doi:10.1016/j.ejmg.2012.10.004

14. Barakeh D, Faqeih E, Anazi S, et al. The many faces of KIF7. Human genome variation. 2015;2:15006. doi:10.1038/hgv.2015.6

15. Ibisler A, Hehr U, Barth A, Koch M, Epplen JT, Hoffjan S. Novel KIF7 Mutation in a Tunisian Boy with Acrocallosal Syndrome: Case Report and Review of the Literature. Molecular syndromology. 2015;6(4):173–180. doi:10.1159/000439414

16. Tunovic S, Baranano KW, Barkovich JA, Strober JB, Jamal L, Slavotinek AM. Novel KIF7 missense substitutions in two patients presenting with multiple malformations and features of acrocallosal syndrome. American journal of medical genetics Part A. 2015;167A(11):2767–2776. doi:10.1002/ajmg.a.37249

17. Asadollahi R, Strauss JE, Zenker M, et al. Clinical and experimental evidence suggest a link between KIF7 and C5orf42-related ciliopathies through Sonic Hedgehog signaling. European journal of human genetics : EJHG. 2018;26(2):197–209. doi:10.1038/s41431-017-0019-9

18. Niceta M, Dentici ML, Ciolfi A, et al. Co-occurrence of mutations in KIF7 and KIAA0556 in Joubert syndrome with ocular coloboma, pituitary malformation and growth hormone deficiency: a case report and literature review. BMC pediatrics. 2020;20(1):120. doi:10.1186/s12887-020-2019-0

19. Dafinger C, Liebau MC, Elsayed SM, et al. Mutations in KIF7 link Joubert syndrome with Sonic Hedgehog signaling and microtubule dynamics. The Journal of clinical investigation. 2011;121(7):2662–2667. doi:10.1172/JCI43639

20. Chiang C, Litingtung Y, Lee E, et al. Cyclopia and defective axial patterning in mice lacking Sonic hedgehog gene function. Nature. 1996;383(6599):407–413. doi:10.1038/383407a0

21. Shimamura K, Rubenstein JLR. Inductive interactions direct early regionalization of the mouse forebrain. Development. 1997;124(14):2709–2718. doi:10.1242/dev.124.14.2709

22. Kohtz JD, Baker DP, Corte G, Fishell G. Regionalization within the mammalian telencephalon is mediated by changes in responsiveness to Sonic Hedgehog. Development. 1998;125(24):5079–5089. doi:10.1242/dev.125.24.5079

23. Machold R, Hayashi S, Rutlin M, et al. Sonic Hedgehog Is Required for Progenitor Cell Maintenance in Telencephalic Stem Cell Niches. Neuron. 2003;39(6):937–950. doi:10.1016/S0896-6273(03)00561-0

24. Xu Q, Wonders CP, Anderson SA. Sonic hedgehog maintains the identity of cortical interneuron progenitors in the ventral telencephalon. Development. 2005;132(22):4987–4998. doi:10.1242/dev.02090

25. Xu Q, Guo L, Moore H, Waclaw RR, Campbell K, Anderson SA. Sonic Hedgehog Signaling Confers Ventral Telencephalic Progenitors with Distinct Cortical Interneuron Fates. Neuron. 2010;65(3):328–340. doi:10.1016/j.neuron.2010.01.004

26. Sousa VH, Fishell G. Sonic hedgehog functions through dynamic changes in temporal competence in the developing forebrain. Current Opinion in Genetics & Development. 2010;20(4):391–399. doi:10.1016/j.gde.2010.04.008

27. Komada M, Saitsu H, Kinboshi M, Miura T, Shiota K, Ishibashi M. Hedgehog signaling is involved in development of the neocortex. Development. 2008;135(16):2717–2727. doi:10.1242/dev.015891

28. Dahmane N, Sánchez P, Gitton Y, et al. The Sonic Hedgehog-Gli pathway regulates dorsal brain growth and tumorigenesis. Development. 2001;128(24):5201–5212. doi:10.1242/dev.128.24.5201

29. Baudoin JP, Viou L, Launay PS, et al. Tangentially migrating neurons assemble a primary cilium that promotes their reorientation to the cortical plate. Neuron. 2012;76(6):1108–1122. doi:10.1016/j.neuron.2012.10.027

30. Higginbotham H, Eom TY, Mariani LE, et al. Arl13b in primary cilia regulates the migration and placement of interneurons in the developing cerebral cortex. Developmental cell. 2012;23(5):925–938. doi:10.1016/j.devcel.2012.09.019

31. Komada M. Sonic hedgehog signaling coordinates the proliferation and differentiation of neural stem/progenitor cells by regulating cell cycle kinetics during development of the neocortex. Congenital anomalies. 2012;52(2):72–77. doi:10.1111/j.1741-4520.2012.00368.x

32. Komada M, Iguchi T, Takeda T, Ishibashi M, Sato M. Smoothened controls cyclin D2 expression and regulates the generation of intermediate progenitors in the developing cortex. Neuroscience letters. 2013;547:87–91. doi:10.1016/j.neulet.2013.05.006

33. Hasenpusch-Theil K, Theil T. The Multifaceted Roles of Primary Cilia in the Development of the Cerebral Cortex. Frontiers in cell and developmental biology. 2021;9:630161. doi:10.3389/fcell.2021.630161

34. Hui CC, Slusarski D, Platt KA, Holmgren R, Joyner AL. Expression of Three Mouse Homologs of the Drosophila Segment Polarity Gene cubitus interruptus, Gli, Gli-2, and Gli-3, in Ectoderm- and Mesoderm-Derived Tissues Suggests Multiple Roles during Postimplantation Development. Developmental Biology. 1994;162(2):402–413. doi:10.1006/dbio.1994.1097

35. Yu W, Wang Y, McDonnell K, Stephen D, Bai CB. Patterning of ventral telencephalon requires positive function of Gli transcription factors. Developmental Biology. 2009;334(1):264–275. doi:10.1016/j.ydbio.2009.07.026

36. Park HL, Bai C, Platt KA, et al. Mouse *Gli1* mutants are viable but have defects in SHH signaling in combination with a *Gli2* mutation. Development. 2000;127(8):1593–1605. doi:10.1242/dev.127.8.1593

37. Bai CB, Joyner AL. *Gli1* can rescue the in vivo function of *Gli2*. Development. 2001;128(24):5161–5172. doi:10.1242/dev.128.24.5161

38. Ding Q, Motoyama J, Gasca S, et al. Diminished Sonic hedgehog signaling and lack of floor plate differentiation in *Gli2* mutant mice. Development. 1998;125(14):2533–2543. doi:10.1242/dev.125.14.2533

39. Tole S, Ragsdale CW, Grove EA. Dorsoventral Patterning of the Telencephalon Is Disrupted in the Mouse Mutant extra-toesJ. Developmental Biology. 2000;217(2):254–265. doi:10.1006/dbio.1999.9509

40. Rallu M, Machold R, Gaiano N, Corbin JG, McMahon AP, Fishell G. Dorsoventral patterning is established in the telencephalon of mutants lacking both Gli3 and Hedgehog signaling. Development. 2002;129(21):4963–4974. doi:10.1242/dev.129.21.4963

41. Wang H, Ge G, Uchida Y, Luu B, Ahn S. *Gli3* Is Required for Maintenance and Fate Specification of Cortical Progenitors. J Neurosci. 2011;31(17):6440–6448. doi:10.1523/JNEUROSCI.4892-10.2011

42. Wilson SL, Wilson JP, Wang C, Wang B, McConnell SK. Primary cilia and Gli3 activity regulate cerebral cortical size. Developmental Neurobiology. 2012;72(9):1196–1212. doi:10.1002/dneu.20985

43. Hasenpusch-Theil K, West S, Kelman A, et al. Gli3 controls the onset of cortical neurogenesis by regulating the radial glial cell cycle through *Cdk6* expression. Development. 2018;145(17):dev163147. doi:10.1242/dev.163147

44. Putoux A, Baas D, Paschaki M, et al. Altered GLI3 and FGF8 signaling underlies acrocallosal syndrome phenotypes in Kif7 depleted mice. Human molecular genetics. 2019;28(6):877–887. doi:10.1093/hmg/ddy392

45. Guo J, Higginbotham H, Li J, et al. Developmental disruptions underlying brain abnormalities in ciliopathies. Nature communications. 2015;6:7857. doi:10.1038/ncomms8857

46. Maurya AK, Ben J, Zhao Z, et al. Positive and negative regulation of Gli activity by Kif7 in the zebrafish embryo. PLoS genetics. 2013;9(12):e1003955. doi:10.1371/journal.pgen.1003955

47. Klejnot M, Kozielski F. Structural insights into human Kif7, a kinesin involved in Hedgehog signalling. *Acta crystallographica Section D*, Biological crystallography. 2012;68(Pt 2):154–159. doi:10.1107/S0907444911053042

48. Chen J, Laclef C, Moncayo A, et al. The ciliopathy gene Rpgrip1l is essential for hair follicle development. The Journal of investigative dermatology. 2015;135(3):701–709. doi:10.1038/jid.2014.483

49. Walczak-Sztulpa J, Eggenschwiler J, Osborn D, et al. Cranioectodermal Dysplasia, Sensenbrenner syndrome, is a ciliopathy caused by mutations in the IFT122 gene. American journal of human genetics. 2010;86(6):949–956. doi:10.1016/j.ajhg.2010.04.012

50. Price DJ, Kennedy H, Dehay C, et al. The development of cortical connections. The European journal of neuroscience. 2006;23(4):910–920. doi:10.1111/j.1460-9568.2006.04620.x

51. Hatanaka Y, Yamauchi K. Excitatory Cortical Neurons with Multipolar Shape Establish Neuronal Polarity by Forming a Tangentially Oriented Axon in the Intermediate Zone. Cerebral Cortex. 2013;23(1):105–113. doi:10.1093/cercor/bhr383

52. Kon E, Cossard A, Jossin Y. Neuronal Polarity in the Embryonic Mammalian Cerebral Cortex. Front Cell Neurosci. 2017;11:163. doi:10.3389/fncel.2017.00163

53. Braisted JE, Catalano SM, Stimac R, et al. Netrin-1 Promotes Thalamic Axon Growth and Is Required for Proper Development of the Thalamocortical Projection. J Neurosci. 2000;20(15):5792–5801. doi:10.1523/JNEUROSCI.20-15-05792.2000

54. Magnani D, Morlé L, Hasenpusch-Theil K, et al. The ciliogenic transcription factor Rfx3 is required for the formation of the thalamocortical tract by regulating the patterning of prethalamus and ventral telencephalon. Human Molecular Genetics. 2015;24(9):2578–2593. doi:10.1093/hmg/ddv021

55. Métin C, Godement P. The Ganglionic Eminence May Be an Intermediate Target for Corticofugal and Thalamocortical Axons. J Neurosci. 1996;16(10):3219–3235. doi:10.1523/JNEUROSCI.16-10-03219.1996

56. Denaxa M, Chan CH, Schachner M, Parnavelas JG, Karagogeos D. The adhesion molecule TAG-1 mediates the migration of cortical interneurons from the ganglionic eminence along the corticofugal fiber system. Development. 2001;128(22):4635–4644. doi:10.1242/dev.128.22.4635

57. Tanaka D, Nakaya Y, Yanagawa Y, Obata K, Murakami F. Multimodal tangential migration of neocortical GABAergic neurons independent of GPI-anchored proteins. Development. 2003;130(23):5803–5813. doi:10.1242/dev.00825

58. Yokota Y, Ghashghaei HT, Han C, Watson H, Campbell KJ, Anton ES. Radial Glial Dependent and Independent Dynamics of Interneuronal Migration in the Developing Cerebral Cortex. Chan-Ling T, ed. PLoS ONE. 2007;2(8):e794. doi:10.1371/journal.pone.0000794

59. Tiveron MC, Rossel M, Moepps B, et al. Molecular interaction between projection neuron precursors and invading interneurons via stromal-derived factor 1 (CXCL12)/CXCR4 signaling in the cortical subventricular zone/intermediate zone. The Journal of neuroscience : the official journal of the Society for Neuroscience. 2006;26(51):13273–13278. doi:10.1523/JNEUROSCI.4162-06.2006

60. Stumm RK, Zhou C, Ara T, et al. CXCR4 Regulates Interneuron Migration in the Developing Neocortex. J Neurosci. 2003;23(12):5123–5130. doi:10.1523/JNEUROSCI.23-12-05123.2003

61. Li G, Adesnik H, Li J, et al. Regional Distribution of Cortical Interneurons and Development of Inhibitory Tone Are Regulated by Cxcl12/Cxcr4 Signaling. J Neurosci. 2008;28(5):1085–1098. doi:10.1523/JNEUROSCI.4602-07.2008

62. Atkins M, Wurmser M, Darmon M, Roche F, Nicol X, Métin C. CXCL12 targets the primary cilium cAMP/cGMP ratio to regulate cell polarity during migration. Nat Commun. 2023;14(1):8003. doi:10.1038/s41467-023-43645-w

63. Caronia-Brown G, Grove EA. Timing of Cortical Interneuron Migration Is Influenced by the Cortical Hem. Cerebral Cortex. 2011;21(4):748–755. doi:10.1093/cercor/bhq142

64. Bellion A. Nucleokinesis in Tangentially Migrating Neurons Comprises Two Alternating Phases: Forward Migration of the Golgi/Centrosome Associated with Centrosome Splitting and Myosin Contraction at the Rear. Journal of Neuroscience. 2005;25(24):5691–5699. doi:10.1523/JNEUROSCI.1030-05.2005

65. Sahara S, Kawakami Y, Izpisua Belmonte JC, O’Leary DD. Sp8 exhibits reciprocal induction with Fgf8 but has an opposing effect on anterior-posterior cortical area patterning. Neural development. 2007;2:10. doi:10.1186/1749-8104-2-10

66. Nery S, Wichterle H, Fishell G. Sonic hedgehog contributes to oligodendrocyte specification in the mammalian forebrain. Development. 2001;128(4):527–540. doi:10.1242/dev.128.4.527

67. Lee J, Ekker S, von Kessler D, Porter J, Sun B, Beachy P. Autoproteolysis in hedgehog protein biogenesis. Science. 1994;266(5190):1528–1537. doi:10.1126/science.7985023

68. Loulier K, Ruat M, Traiffort E. Analysis of hedgehog interacting protein in the brain and its expression in nitric oxide synthase-positive cells. Neuroreport. 2005;16(17):1959–1962. doi:10.1097/01.wnr.0000187632.91375.81

69. Kiecker C, Lumsden A. Hedgehog signaling from the ZLI regulates diencephalic regional identity. Nat Neurosci. 2004;7(11):1242–1249. doi:10.1038/nn1338

70. Huang X, Liu J, Ketova T, et al. Transventricular delivery of Sonic hedgehog is essential to cerebellar ventricular zone development. Proc Natl Acad Sci USA. 2010;107(18):8422–8427. doi:10.1073/pnas.0911838107

71. Theil T. Gli3 is required for the specification and differentiation of preplate neurons. Developmental Biology. 2005;286(2):559–571. doi:10.1016/j.ydbio.2005.08.033

72. He M, Subramanian R, Bangs F, et al. The kinesin-4 protein Kif7 regulates mammalian Hedgehog signalling by organizing the cilium tip compartment. Nature cell biology. 2014;16(7):663–672. doi:10.1038/ncb2988

73. Lai FPL, Li Z, Zhou T, et al. Ciliary protein Kif7 regulates Gli and Ezh2 for initiating the neuronal differentiation of enteric neural crest cells during development. Sci Adv. 2021;7(42):eabf7472. doi:10.1126/sciadv.abf7472

74. Marín O. Interneuron dysfunction in psychiatric disorders. Nat Rev Neurosci. 2012;13(2):107–120. doi:10.1038/nrn3155

75. Barkovich AJ, Dobyns WB, Guerrini R. Malformations of Cortical Development and Epilepsy. Cold Spring Harbor Perspectives in Medicine. 2015;5(5):a022392–a022392. doi:10.1101/cshperspect.a022392

76. Yue Y, Engelke MF, Blasius TL, Verhey KJ. Hedgehog-induced ciliary trafficking of kinesin-4 motor KIF7 requires intraflagellar transport but not KIF7’s microtubule binding. Molecular biology of the cell. 2022;33(1):br1. doi:10.1091/mbc.E21-04-0215

77. Willaredt MA, Hasenpusch-Theil K, Gardner HAR, et al. A Crucial Role for Primary Cilia in Cortical Morphogenesis. J Neurosci. 2008;28(48):12887–12900. doi:10.1523/JNEUROSCI.2084-08.2008

78. Magnani D, Hasenpusch-Theil K, Jacobs EC, Campagnoni AT, Price DJ, Theil T. The Gli3 hypomorphic mutation Pdn causes selective impairment in the growth, patterning, and axon guidance capability of the lateral ganglionic eminence. The Journal of neuroscience : the official journal of the Society for Neuroscience. 2010;30(41):13883–13894. doi:10.1523/JNEUROSCI.3650-10.2010

79. Magnani D, Hasenpusch-Theil K, Theil T. Gli3 controls subplate formation and growth of cortical axons. Cereb Cortex. 2013;23(11):2542–2551. doi:10.1093/cercor/bhs237

80. Theil T. Gli3 is required for the specification and differentiation of preplate neurons. Developmental biology. 2005;286(2):559–571. doi:10.1016/j.ydbio.2005.08.033

81. Rash BG, Grove EA. Shh and Gli3 regulate formation of the telencephalic–diencephalic junction and suppress an isthmus-like signaling source in the forebrain. Developmental Biology. 2011;359(2):242–250. doi:10.1016/j.ydbio.2011.08.026

82. Vilboux T, Malicdan MC, Roney JC, et al. CELSR2, encoding a planar cell polarity protein, is a putative gene in Joubert syndrome with cortical heterotopia, microophthalmia, and growth hormone deficiency. American journal of medical genetics Part A. 2017;173(3):661–666. doi:10.1002/ajmg.a.38005

83. Boutin C, Goffinet AM, Tissir F. Celsr1–3 Cadherins in PCP and Brain Development. In: Current Topics in Developmental Biology. Vol 101. Elsevier; 2012:161–183. doi:10.1016/B978-0-12-394592-1.00010-7

84. Hakanen J, Parmentier N, Sommacal L, et al. The Celsr3-Kif2a axis directs neuronal migration in the postnatal brain. Progress in Neurobiology. 2022;208:102177. doi:10.1016/j.pneurobio.2021.102177

85. Alcauter S, Lin W, Smith JK, et al. Development of thalamocortical connectivity during infancy and its cognitive correlations. The Journal of neuroscience : the official journal of the Society for Neuroscience. 2014;34(27):9067–9075. doi:10.1523/JNEUROSCI.0796-14.2014

86. Jakab A, Natalucci G, Koller B, Tuura R, Ruegger C, Hagmann C. Mental development is associated with cortical connectivity of the ventral and nonspecific thalamus of preterm newborns. Brain and behavior. 2020;10(10):e01786. doi:10.1002/brb3.1786

87. Tissir F, Bar I, Jossin Y, De Backer O, Goffinet AM. Protocadherin Celsr3 is crucial in axonal tract development. Nature neuroscience. 2005;8(4):451–457. doi:10.1038/nn1428

88. Wang Y, Thekdi N, Smallwood PM, Macke JP, Nathans J. Frizzled-3 is required for the development of major fiber tracts in the rostral CNS. The Journal of neuroscience : the official journal of the Society for Neuroscience. 2002;22(19):8563–8573. doi:10.1523/JNEUROSCI.22-19-08563.2002

89. Moreau MX, Saillour Y, Cwetsch AW, Pierani A, Causeret F. Single-cell transcriptomics of the early developing mouse cerebral cortex disentangle the spatial and temporal components of neuronal fate acquisition. Development. 2021;148(14):dev197962. doi:10.1242/dev.197962

90. Zakaria M, Ferent J, Hristovska I, et al. The Shh receptor Boc is important for myelin formation and repair. Development. 2019;146(9). doi:10.1242/dev.172502

91. Vyas N, Walvekar A, Tate D, et al. Vertebrate Hedgehog is secreted on two types of extracellular vesicles with different signaling properties. Scientific reports. 2014;4:7357. doi:10.1038/srep07357

92. Eitan E, Petralia RS, Wang YX, Indig FE, Mattson MP, Yao PJ. Probing extracellular Sonic hedgehog in neurons. Biology open. 2016;5(8):1086–1092. doi:10.1242/bio.019422

93. Hasenpusch-Theil K, Laclef C, Colligan M, et al. A transient role of the ciliary gene Inpp5e in controlling direct versus indirect neurogenesis in cortical development. eLife. 2020;9. doi:10.7554/eLife.58162

94. Higginbotham H, Guo J, Yokota Y, et al. Arl13b-regulated cilia activities are essential for polarized radial glial scaffold formation. Nature neuroscience. 2013;16(8):1000–1007. doi:10.1038/nn.3451

95. Han YG, Spassky N, Romaguera-Ros M, et al. Hedgehog signaling and primary cilia are required for the formation of adult neural stem cells. Nat Neurosci. 2008;11(3):277–284. doi:10.1038/nn2059

96. Lysko DE, Putt M, Golden JA. SDF1 regulates leading process branching and speed of migrating interneurons. The Journal of neuroscience : the official journal of the Society for Neuroscience. 2011;31(5):1739–1745. doi:10.1523/JNEUROSCI.3118-10.2011

97. Tiveron MC, Hirsch MR, Brunet JF. The Expression Pattern of the Transcription Factor Phox2 Delineates Synaptic Pathways of the Autonomic Nervous System. J Neurosci. 1996;16(23):7649–7660. doi:10.1523/JNEUROSCI.16-23-07649.1996

98. Renier N, Adams EL, Kirst C, et al. Mapping of Brain Activity by Automated Volume Analysis of Immediate Early Genes. Cell. 2016;165(7):1789–1802. doi:10.1016/j.cell.2016.05.007

99. Gorelik R, Gautreau A. Quantitative and unbiased analysis of directional persistence in cell migration. Nat Protoc. 2014;9(8):1931–1943. doi:10.1038/nprot.2014.131

